# The Conserved *Colletotrichum* spp. Effector CEC3 Induces Nuclear Expansion and Cell Death in Plants

**DOI:** 10.1101/2021.03.18.435704

**Authors:** Ayako Tsushima, Mari Narusaka, Pamela Gan, Naoyoshi Kumakura, Ryoko Hiroyama, Naoki Kato, Shunji Takahashi, Yoshitaka Takano, Yoshihiro Narusaka, Ken Shirasu

**Affiliations:** Graduate School of Science, The University of Tokyo, Bunkyo, Japan; Center for Sustainable Resource Science, RIKEN, Yokohama, Japan; Research Institute for Biological Sciences Okayama, Kaga-gun, Japan; Center for Sustainable Resource Science, RIKEN, Wako, Japan; Graduate School of Agriculture, Kyoto University, Kyoto, Japan

**Keywords:** core effector, comparative genomics, nuclear expansion, cell death, *Colletotrichum*, fungal plant pathogen

## Abstract

Plant pathogens secrete small proteins, known as effectors, that promote infection by manipulating host cells. Members of the phytopathogenic fungal genus *Colletotrichum* collectively have a broad host range and generally adopt a hemibiotrophic lifestyle that includes an initial biotrophic phase and a later necrotrophic phase. We hypothesized that *Colletotrichum* fungi use a set of conserved effectors during infection to support the two phases of their hemibiotrophic lifestyle. This study aimed to examine this hypothesis by identifying and characterizing conserved effectors among *Colletotrichum* fungi. Comparative genomic analyses using genomes of ascomycete fungi with different lifestyles identified seven effector candidates that are conserved across the genus *Colletotrichum*. Transient expression assays showed that one of these conserved effectors, CEC3, induces nuclear expansion and cell death in *Nicotiana benthamiana*, suggesting that CEC3 is involved in promoting host cell death during infection. Nuclear expansion and cell death induction were commonly observed in CEC3 homologs from four different *Colletotrichum* species that vary in host specificity. Thus, CEC3 proteins could represent a novel class of core effectors with functional conservation in the genus C*olletotrichum*.

## Introduction

Plant pathogens have adopted different strategies to extract nutrients from their individual hosts: evading or disabling the host immune system to establish a parasitic relationship with living cells (biotrophy), induction of a lethal response (necrotrophy), or by using a combination of these strategies (hemibiotrophy). These pathogens have evolved an array of secreted proteins that manipulate host cell responses, collectively referred to as effectors, that allow them to establish a defined relationship with their hosts. Although effectors play pivotal roles in establishing parasitic interactions, some effectors are also detected by plants via immune receptors encoded by resistance (R) genes, thereby triggering strong host immune responses (Dodds and Rathjen, 2010). The genes that encode effectors are called ‘avirulence genes’ because the plant response triggered by recognition of effectors by cognate immune receptors results in the loss of virulence. Thus, effectors both positively and negatively impact the ability of a pathogen to establish a disease state, depending on the host genotype. The importance of host and pathogen genotypes in the outcome of infection is illustrated by the zig-zag model wherein pathogens continuously evolve new effectors to overcome plant defense responses and hosts evolve receptors that recognize the newly evolved effectors, resulting in disease resistance (Jones and Dangl, 2006). As a corollary to this model, the ability to infect a host that can perceive a particular effector requires that the pathogen lose or alter the effector to escape recognition. Consistent with this model, previous studies have shown that known avirulence effectors often lack homologs in closely-related lineages as a result of high selection pressure in the arms race between host and pathogen (Sanchez-Vallet et al., 2018). In extreme cases, avirulence effector genes such as *Avr4E* and *AvrStb6*, which were isolated from fungal plant pathogens *Cladosporium fulvum* and *Zymoseptoria tritici*, respectively, have only been found in specific strains within a single species (Westerink et al., 2004; Zhong et al., 2017). In contrast, some effectors are widely conserved among different taxa and are required for full virulence on a range of different hosts. For example, many fungal plant pathogens express LysM effectors, which protect fungal cells from plant chitinases and dampen host immune responses (Akcapinar et al., 2015). NIS1 and its homologs are also common among the Ascomycota and Basidiomycota. NIS1 suppresses the kinase activities of BAK1 and BIK1, which are critical for transmitting host immune signaling (Irieda et al., 2019). As another example, Pep1 and its homologs, which inhibit plant peroxidases required for accumulation of reactive oxygen species, are conserved within the fungal order Ustilaginales (Hemetsberger et al., 2012, 2015). Importantly, these conserved effectors contribute to pathogenicity by targeting host proteins that are conserved in a wide range of plant taxa.

*Colletotrichum* is one of the most economically important genera among plant pathogenic fungi because of its ubiquity and ability to cause serious crop losses (Dean et al., 2012). *Colletotrichum* spp. can be grouped into several major monophyletic clades that are termed species complexes (Cannon et al., 2012). Among the species complexes, members of the *Colletotrichum gloeosporioides* species complex tend to have a wide host range as post-harvest pathogens. For example, *Colletotrichum fructicola* infects a wide range of fruits, including strawberry (*Fragaria* × *ananassa*), apple (*Malus domestica*), and avocado (*Persea americana*) (Weir et al., 2012). In contrast, members of other species complexes tend to have more limited host ranges. The *Colletotrichum graminicola* species complex has members that are restricted to infecting gramineous plants, such as *C. graminicola*, which is associated with maize (Crouch and Beirn, 2009). *Colletotrichum higginsianum*, a member of the *Colletotrichum destructivum* species complex, infects Brassicaceae plants, including *Arabidopsis thaliana* (O’Connell et al., 2004). Similarly, *Colletotrichum orbiculare* from the *C. orbiculare* species complex infects Cucurbitaceae plants as well as *Nicotiana benthamiana* (Shen et al., 2001). Therefore, while members of this genus have a collective wide host range, individual *Colletotrichum* spp. host ranges are often much more limited. Despite the host range of each *Colletotrichum* sp., the majority have adapted a hemibiotrophic lifestyle. They develop bulbous primary hyphae within living host cells during the initial biotrophic phase, then induce host death in the subsequent necrotrophic phase, which is characterized by the production of filamentous secondary hyphae (Perfect et al., 1999).

Based on what is known about the genus *Colletotrichum*, we hypothesize that effectors in *Colletotrichum* spp. fall into two classes based on their conservation patterns: 1) specialized effectors, which have recently evolved for adaptation to specific host niches, and 2) conserved effectors, which are generally required for infection of a wide range of plants. To date, more than 100 genomes of *Colletotrichum* spp. have been sequenced due to their agricultural importance and scientific interest (O’Connell et al., 2012; Baroncelli et al., 2014, 2016a; Gan et al., 2016, 2019, 2020; Hacquard et al., 2016). Using these abundant genome resources, it is now feasible to conduct comparative genomic analysis and reverse genetics of *Colletotrichum* spp. Since *Colletotrichum* spp. have a collective broad host range, their effectors conserved within the genus should have important roles during infection across a wide range of host plants. Here, we identified effector candidates and their conservation patterns across ascomycetes with different lifestyles. Among them, the effector candidate ChCEC3 (core effector of *Colletotrichum* fungi 3 from *C. higginsianum*), can induce nuclear expansion and cell death when expressed in *N. benthamiana*. CEC3 homologs from four different *Colletotrichum* species that have different host specificities also induce nuclear expansion and cell death, indicating that their functional role is conserved in the genus *Colletotrichum*.

## Materials and Methods

### Prediction of Effector Candidates

In this study, effector candidates were defined as predicted secreted proteins (*i.e.,* those with a signal peptide sequence but no transmembrane domain) less than 300 amino acids long. SignalP 4.1 (Petersen et al., 2011) and TMHMM 2.0 (Krogh et al., 2001) were used with default settings to predict signal peptides and transmembrane domains, respectively.

### Conservation Patterns of Effector Candidates

Twenty-four ascomycetes that are associated with saprophyte, plant pathogen, or insect pathogen lifestyles, were selected to assess the conservation patterns of the protein sequences (Supplementary Table 1). To identify orthogroups, OrthoFinder v2.2.7 (Emms and Kelly, 2015) was used with default settings. Analyses of all proteins and effector candidates were independently performed. The conservation patterns of CEC proteins were further investigated by performing BLASTP against the NCBI non-redundant protein database (last accessed on 29 June 2020) using ChCECs as the query amino acid sequences with a cutoff *E*-value = 10^−30^ Based on this result, we selected 70 proteomes, including all of the publicly available proteomes of 35 *Colletotrichum* strains and 35 fungal proteomes representing different branches of the Ascomycota (Supplementary Table 2). Then, the amino acid sequences of ChCECs were used as queries for BLASTP against the database generated using the 70 proteomes (cutoff *E*-value = 10^−30^). To identify functional protein domains of CEC3 proteins, InterProScan 5.39-77.0 (Mitchell et al., 2019) was used with default settings. Amino acid sequence alignments and a phylogenetic tree of CEC3 homologs were generated using CLC Genomics Workbench8 (QIAGEN bioinformatics).

### Phylogenetic Analyses

A phylogenetic tree of 24 ascomycetes was generated from the combined alignments of single-copy orthologs conserved in all 24 ascomycetes identified using OrthoFinder v2.2.7 with default settings. Protein sequences were aligned using MAFFT version 7.215 (Katoh et al., 2002) with the --auto settings and trimmed using trimAL v1.2 (Capella-Gutierrez et al., 2009) with the automated1 settings. The concatenated and trimmed alignments were then used to estimate a maximum-likelihood species phylogeny with RAxML version 8.2.11 (Stamatakis, 2014) with 1,000 bootstrap replicates. To generate the maximum-likelihood tree, the PROTGAMMAAUTO setting was used to find the best protein substitution model and the autoMRE setting was used to determine the appropriate number of bootstrap samples. The tree was visualized using iTOL version 4.1 (Letunic and Bork, 2016). A phylogenetic tree of 70 ascomycetes was generated in the same way using the combined alignments of single-copy orthologs conserved across all proteomes (Supplementary Table 2). *Saccharomyces cerevisiae* sequences were used as the outgroup in both trees.

### Cloning

Total RNA was extracted from *C. higginsianum* MAFF 305635, *C. orbiculare* MAFF 240422, and *C. graminicola* MAFF 244463 cultured in potato dextrose (PD) broth (BD Biosciences) at 24°C in the dark for two days. Total RNA was extracted from strawberry (Sachinoka) leaves three days after inoculation with *C. fructicola* Nara gc5 (JCM 39093) as previously described (Gan et al., 2020). RNA was extracted using RNeasy Plant Mini Kit (Qiagen) with DNase I treatment according to the manufacturer’s introductions, and reverse transcribed using ReverTraAce qPCR RT Kit (Toyobo, Co., Ltd.) or SuperScript III Reverse Transcriptase (Thermo Fisher Scientific). cDNAs of *ChCEC2-1*, *ChCEC2-2*, and *ChCEC6* were amplified using primers listed in Supplementary Table 3 and Phusion^®^ High-Fidelity DNA Polymerase (New England Biolabs), then cloned into pCR8/GW/TOPO (Thermo Fisher Scientific). The CDS of *ChCEC3* was synthesized in pDONR/Zeo (Thermo Fisher Scientific) by Invitrogen. The *ChCEC* sequences were transferred into Gateway-compatible pSfinx (pSfinx-GW) (Narusaka et al., 2013) using Gateway LR Clonase Enzyme mix (Thermo Fisher Scientific). cDNAs of *CEC3* without stop codons and the regions encoding predicted signal peptides and stop codons were also amplified and cloned into pCR8/GW/TOPO. Each *CEC3* derivative was transferred into pGWB5 (Nakagawa et al., 2007) with Gateway LR Clonase Enzyme mix. *ChCEC3ΔSP* and *YFP* cloned from pGWB41 (Nakagawa et al., 2007) in pCR8/GW/TOPO were also transferred into pEDV6 (Fabro et al., 2011) using Gateway LR Clonase Enzyme mix. We deposited the cDNA sequences of *CoCEC3-2.2* from *C. orbiculare* MAFF 240422 and *CgCEC3* from *C. graminicola* MAFF 244463 in NCBI GenBank under the accession numbers MW528236 and MW528237, respectively.

To create transformation vectors for overexpression or knock-out mutations, we first generated pAGM4723_TEF_GFP_scd1_HygR using Golden Gate cloning (Engler et al., 2014) (Supplementary Figure 1). To generate pAGM4723_TEF_ChCEC3g_scd1_HygR, genomic DNA encoding *ChCEC3* and the linearized pAGM4723_TEF_GFP_scd1_HygR lacking the GFP sequence were amplified using KOD -Plus- Neo (Toyobo, Co., Ltd.), then the fragments were circularized using In-Fusion HD (Takara Bio Inc.). To generate pAGM4723-ChCEC3KO, 5’ and 3’ 2 kb genomic fragments of *ChCEC3*, the hygromycin resistance cassette, and linearized pAGM4723 were amplified using KOD-Plus- Neo. These fragments were circularized using In-Fusion HD. Genomic DNA of *C. higginsianum* MAFF 305635 was extracted using DNeasy Plant Mini Kit (Qiagen).

### Cell Death-Inducing Effector Candidate Screening

*Agrobacterium tumefaciens* strain GV3101 was used to screen for cell death-inducing effector candidates. Binary vectors were transformed into *A. tumefaciens* with the freeze-thaw method or by electroporation. After transformation, *A. tumefaciens* was cultured on Luria-Bertani (LB) agar (Merck KGaA) containing 100 μg/ml rifampicin and 50 μg/ml kanamycin at 28°C for two days. *A. tumefaciens* transformant colonies were purified and cultured in LB broth supplemented with 100 μg/ml rifampicin and 50 μg/ml kanamycin at 28°C for two days with shaking at 120 rpm for agroinfiltration. Bacterial cells were collected by centrifugation and resuspended in 10 mM MgCl_2_, 10 mM MES (pH 5.6), and 150 μM acetosyringone. Each bacterial suspension was adjusted to OD600 = 0.3. Suspensions were infiltrated into 4-week-old *N. benthamiana* leaves grown at 25°C under long-day conditions (16 h light/8 h dark) using 1 ml needleless syringes. Plant cell death was visualized six days after infiltration under UV illumination.

### Cell Death Assays

Binary vectors were transformed into *A. tumefaciens* strain AGL1 with the freeze-thaw method or by electroporation. We used *A. tumefaciens* strain C58C1 pCH32 harboring pBCKH 35S promoter::GFP as a negative control to express 35S-driven GFP (Mitsuda et al., 2006). *A. tumefaciens* cultures were prepared as described above. For cell death assays, bacterial suspensions were adjusted to OD600 = 0.5. Suspensions were infiltrated into 4-week-old *N. benthamiana* leaves using 1 ml needleless syringes. Plant cell death was visualized by trypan blue staining five days after infiltration: each *N. benthamiana* leaf was boiled in 20 ml of alcoholic lactophenol (ethanol: phenol: glycerol: lactic acid: water (4: 1: 1: 1: 1, v/v/v/v/v)) containing 0.1 μg/ml trypan blue for 15 minutes and left overnight at room temperature. Boiled leaves were destained with 40% chloral hydrate solution for three to five days before being photographed. Eight different infiltrated leaves were observed for each construct.

### Confocal Microscopy

*A. tumefaciens* strain AGL1 prepared as above (OD600 = 0.3) and carrying binary vectors was infiltrated into 4-week-old *N. benthamiana* leaves. Protein localization was assessed 24 or 36 hours after infiltration in epidermal cells of *N. benthamiana* using Leica SP8 (Leica Microsystems) or Zeiss LSM 700 (Carl Zeiss AG) microscopes. For DAPI (4′,6-diamidino-2-phenylindole) staining, the Staining Buffer in CyStain UV precise P (Sysmex America, Inc.) was infiltrated into *N. benthamiana* leaves using 1 ml needleless syringes one hour before observation. To image GFP fluorescence, excitation was at 488 nm and emission was collected between 495 and 550 nm. For mCherry fluorescence, excitation was at 555 nm and emission was between 505 and 600 nm. DAPI fluorescence was excited at 405 nm and observed between 410 and 480 nm. Chlorophyll autofluorescence was excited at 633 nm and observed between 638 and 700 nm. Nuclear diameters were measured at their narrowest points (minor axes) using ImageJ 1.51k (Schneider et al., 2012).

### Transformation and Infection of *C. higginsianum*

*C. higginsianum* transformants were obtained using *A. tumefaciens* as described in Supplementary Material 1. Transformants were genotyped by PCR using primers listed in Supplementary Table 3. We confirmed constitutive expression by semi-quantitative PCR in fungal hyphae cultured in PD broth for two days at 24°C in the dark for *ChCEC3* over-expressing lines. For lesion area measurement assays, *Arabidopsis thaliana* Col-0 plants were grown at 22°C with a 10-h photoperiod for four weeks. *C. higginsianum* MAFF 305635 and the transformants were cultured on PDA at 24°C under 12-h black-light blue fluorescent bulb light/12-h dark conditions for one week. Lesion area measurement assays were performed as described (Tsushima et al. 2019a). Three leaves per plant were inoculated with 5-μl droplets of conidial suspensions at 5 × 10^5^ conidia/ml. Symptoms were observed six days after inoculation, and lesion areas were measured using the color threshold function of ImageJ 1.51k (Schneider et al., 2012) using the following settings: hue, 0-255; saturation, 110-140; and brightness, 0-255 with a square region of interest. For RT-qPCR analysis, we used fungal hyphae cultured in PD broth for two days at 24°C in the dark as *in vitro* samples and epidermal tissues from infected leaves as *in planta* samples. Epidermal tissues were sampled following the methods described by Takahara et al., 2009 and Kleemann et al., 2012. Approximately 100 detached four-week-old *A. thaliana* leaves per sample were placed on a piece of wet paper towel in a plastic dish. The abaxial leaf surface was inoculated with approximately 50 μl of conidial suspension at 5 × 10^6^ conidia/ml using a micropipette. After inoculation, the lid of the plastic dish was secured using Parafilm to maintain 100% humidity during infection. Inoculated leaves were incubated at 22°C in the dark until sample collection. The epidermis was peeled from the infected abaxial leaf surface using tweezers and double-sided tape, then immediately flash frozen in liquid nitrogen and stored at −80°C until RNA extraction.

### RT-qPCR Analysis

Total RNA was extracted using RNeasy Plant Mini Kit with DNase I treatment according to the manufacturer’s introductions. RT-qPCR was performed using ReverTra Ace (Toyobo, Co., Ltd.) and THUNDERBIRD SYBR qPCR Mix (Toyobo, Co., Ltd.). Reactions were run on an Mx3000P QPCR system and analyzed with MxPro QPCR software (Stratagene California) using primers listed in Supplementary Table 3. To confirm progression of infection at each time point, a few inoculated leaves were stained with 1 ml/leaf alcoholic lactophenol containing 0.1 μg/ml trypan blue for five minutes at 95°C and left overnight at room temperature. Boiled leaves were destained with 40% chloral hydrate solution for three to five days before being observing fungal structures using an Olympus BX51 microscope (Olympus Corporation).

### *Pseudomonas syringae* pv. *tomato* DC3000 Transformation and Infection

Plasmid constructs pEDV6:ChCEC3ΔSP and pEDV6:YFP were mobilized from *Escherichia coli* DH5a into *Pseudomonas syringae* pv. *tomato* (*Pto*) DC3000 by triparental mating using *E. coli* HB101 (pRK2013) as the helper strain. *Pto* DC3000 carrying pEDV6:ChCEC3ΔSP or pEDV6:YFP was cultured on LB agar containing 100 μg/ml rifampicin and 20 μg/ml gentamicin at 28°C in the dark for two days. Bacterial cells collected from LB agar were suspended in 10 mM MgCl_2_, adjusted to OD600 = 0.0002, and infiltrated into three leaves per plant using 1 ml needleless syringes. Leaf tissue was collected using an 8 mm diameter biopsy punch four days after inoculation, and homogenized in 1 ml distilled water. Homogenized tissue was diluted in a tenfold dilution series from 5 × 10^−3^ to 5 × 10^−6^ and spotted onto LB agar containing 100 μg/ml rifampicin. After overnight incubation at 24°C, colony forming units per unit area (cfu)/cm^2^ were determined.

### Immunoblotting

To examine protein expression of CEC3 homologs expressed by agroinfiltration, protein samples were extracted using GTEN-buffer (10% (v/v) Glycerol, 25 mM Tris-HCl (pH 7.5), 1 mM EDTA, 150 mM NaCl, 10 mM DTT, 1xPlant protease inhibitor cocktail (Sigma)). Proteins were separated on Criterion TGX Precast Gels (4-15%) (Bio-Rad Laboratories, Inc.) and electroblotted onto PVDF membranes using a Trans-Blot Turbo Transfer System (Bio-Rad Laboratories, Inc.). Membranes were blocked in TBS-T with 5% skim milk powder at 4°C overnight and incubated in 1:8000 diluted anti-GFP antibody (Ab290; Abcam) in TBS-T with 5% skim milk powder for one hour at room temperature. After washing with TBS-T, membranes were incubated in 1:10000 diluted anti-rabbit IgG (NA934-1ML; GE Healthcare) in TBS-T for one hour at room temperature. Following a final wash with TBS-T, signals were detected using SuperSignal West Femto Maximum Sensitivity Substrate (Thermo Fisher Scientific) and ImageQuant LAS 4010 (GE Healthcare). Proteins on the membrane were visualized by Coomassie Brilliant Blue (CBB) staining. *A. thaliana* leaves infiltrated with 10 mM MgCl_2_ or *Pto* DC3000 carrying pEDV6:ChCEC3ΔSP or pEDV6:YFP (OD_600_ = 2.0) were sampled 24 hours after inoculation to assess expression of ChCEC3ΔSP and YFP proteins in *Pto* DC3000, followed by immunoblotting as described above. Protein concentrations for each sample were measured using Pierce BCA Protein Assay Kit (Thermo Fisher Scientific) and Infinite F200 PRO (Tecan Group Ltd.), and a total of 100 μg of protein was loaded for each sample onto SDS-PAGE gels. Proteins were detected using 1:5000 diluted anti-HA antibody (Anti-HA-Peroxidase, High Affinity (3F10); Roche) in TBS-T.

## Results

### Identification of Effector Candidates Conserved in *Colletotrichum* spp

To identify *Colletotrichum* conserved candidate effectors, we analyzed the proteomes of 24 ascomycetes, including seven *Colletotrichum* species representing the six species complexes and one minor clade. Putative secreted proteins were classified as effector candidates if their lengths were less than 300 amino acids. To investigate the conservation patterns of all proteins and effector candidates from the 24 ascomycetes, orthogroups of these two datasets were independently determined using OrthoFinder (Emms and Kelly, 2015). This analysis identified 15,521 all protein (AP) orthogroups and 990 effector candidate (EC) orthogroups. The proportion of genes belonging to AP orthogroups ranges from 67.4% in *Botrytis cinerea* to 99.2% in *Colletotrichum chlorophyti*, while the proportion of genes belonging to EC orthogroups ranges from 28.4% in *S. cerevisiae* to 98.9% in *C. chlorophyti* (Figure 1A). The percentage of shared orthogroups between each species indicates that effector candidates are less conserved than all proteins (Figure 1B). Among AP orthogroups, 2,424 (15.61%) were found in all ascomycetes tested. In contrast, there were no EC orthogroups that were conserved across all of the ascomycetes tested. This analysis identified seven EC orthogroups (0.71%) that are conserved in all *Colletotrichum* species, but not the other ascomycetes tested (Figure 1C, Supplementary Table 4). We have designated these effector candidates CEC1 (<u>c</u>ore <u>e</u>ffector of *<u>C</u>olletotrichum*) to CEC7. Among the eight predicted CEC proteins from *C. higginsianum* (ChCECs), ChCEC2 has two homologs (ChCEC2-1 and ChCEC2-2) and the others have one homolog. ChCEC2-2 and ChCEC4 were previously identified as ChEC65 and ChEC98, respectively (Kleemann et al., 2012; Robin et al., 2018) (Supplementary Table 5).

**FIGURE 1.**
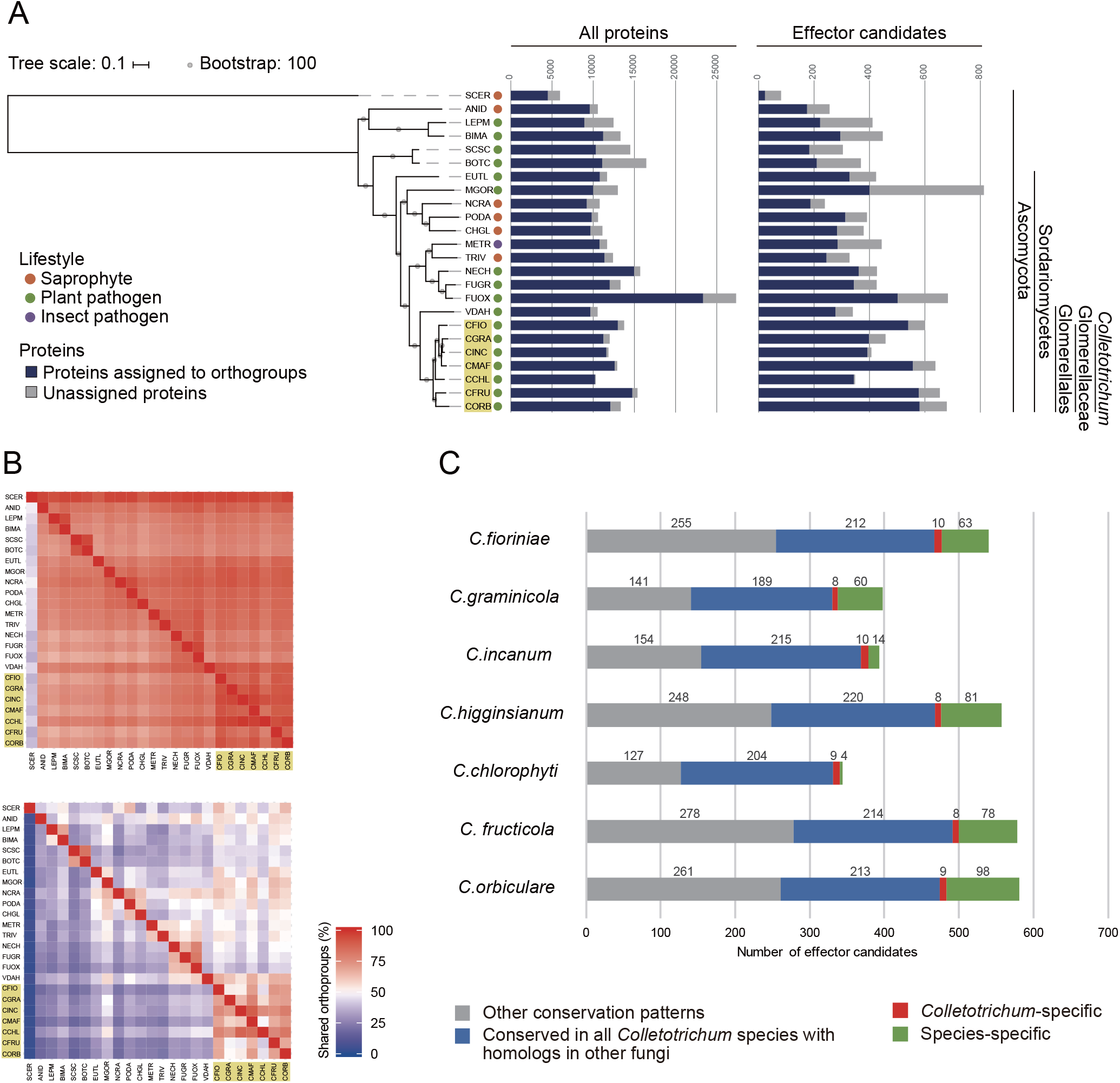
Conservation patterns of proteomes from 24 ascomycete fungi. **(A)** Number of proteins assigned to orthogroups. A maximum-likelihood species phylogeny was drawn based on the alignment of single-copy orthologs obtained using OrthoFinder. Bootstrap values are based on 1,000 replicates. The yellow box indicates *Colletotrichum* species. SCER: *Saccharomyces cerevisiae*, ANID: *Aspergillus nidulans*, LEPM: *Leptosphaeria maculans*, BIMA: *Bipolaris maydis*, SCSC: *Sclerotinia sclerotiorum*, BOTC: *Botrytis cinerea*, EUTL: *Eutypa lata*, MGOR: *Magnaporthe oryzae*, NCRA: *Neurospora crassa*, PODA: *Podospora anserine*, CHGL: *Chaetomium globosum*, METR: *Metarhizium robertsii*, TRIV: *Trichoderma virens*, NECH: *Nectria haematococca*, FUGR: *Fusarium graminearum*, FUOX: *Fusarium oxysporum* f. sp. *lycopersici*, VDAH: *Verticillium dahliae*, CFIO: *C. fioriniae*, CGRA: *C. graminicola*, CINC: *C. incanum*, CMAF: *C. higginsianum*, CCHL: *C. chlorophyti*, CFRU: *C. fructicola*, CORB: *C. orbiculare*. **(B)** Heatmap showing the conservation of orthogroups of all proteins (upper) and effector candidates (lower) between each species. *Colletotrichum* species are highlighted with yellow boxes. **(C)** Conservation patterns of effector candidates from *Colletotrichum* species. The bar chart indicates the number of effector candidates in orthogroups by conservation pattern.

### *CEC3* is Conserved among *Colletotrichum* spp. and the Expression of *ChCEC3* Induces Cell Death in *N. benthamiana*

To assess the conservation of *CEC* genes in greater detail, the amino acid sequences of ChCECs were queried against the NCBI non-redundant protein database (BLASTP, cutoff *E*-value = 10^−30^) (Supplementary Material 2). Based on this result, we selected 70 proteomes, including all publicly available proteomes of 35 *Colletotrichum* strains and 35 fungal proteomes representing different branches of the Ascomycota. The conservation patterns of *CEC* genes were further investigated against the database generated using the 70 proteomes (BLASTP, cutoff *E*-value = 10^−30^) (Figure 2A). This analysis revealed that CEC1, CEC4, and CEC7 are specifically found in *Colletotrichum* spp., but they are not conserved across the genus. In contrast, highly similar homologs of CEC2, CEC3, and CEC6 are conserved across the *Colletotrichum* genus as well as some other ascomycetes. A supplementary analysis to determine if ChCECs have known functional domains (InterProScan 5.39-77.0; Mitchell et al., 2019) indicated that, except for signal peptides, they have no known functional domains (Figure 2B).

**FIGURE 2.**
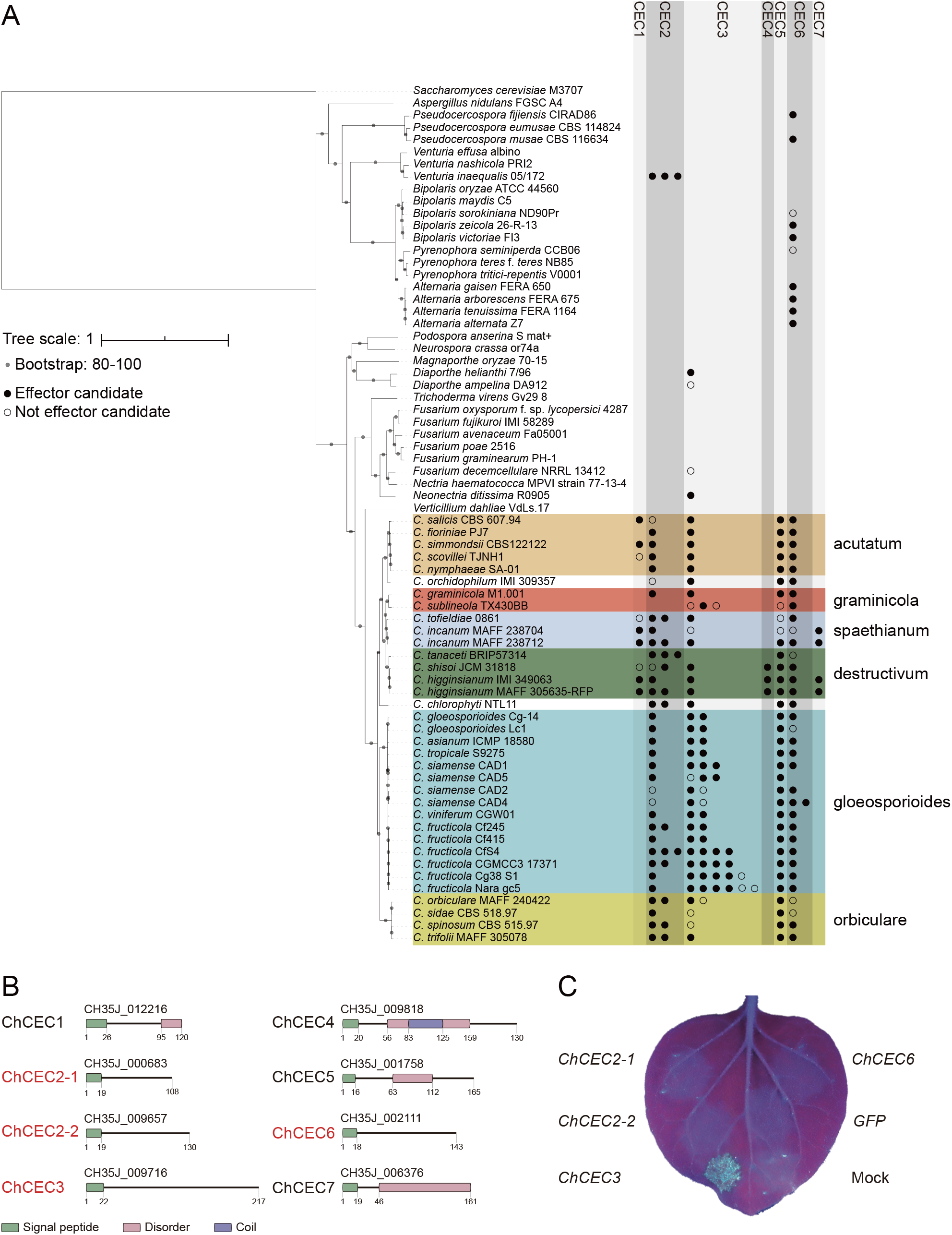
**(A)** Conservation patterns of *CEC* genes examined using BLASTP. The maximum-likelihood species phylogeny was drawn based on the alignment patterns of single-copy orthologs obtained using OrthoFinder. Bootstrap values are based on 1,000 replicates. Colored boxes represent species complexes within the *Colletotrichum* genus. **(B)** Predicted functional domains of CEC proteins found in *C. higginsianum* MAFF 305635-RFP. Red letters indicate effector candidates that are up-regulated during infection as reported previously (O’Connell et al., 2012). **(C)** Representative cell death assay result using *A. tumefaciens* harboring pSfinx vectors. The image was taken six days after infiltration under UV illumination. No GFP fluorescence was visible, as it was weaker than fluorescence from dead leaf tissue.

Since *Colletotrichum* spp. are hemibiotrophic plant pathogens that employ a necrotrophic phase during which host cells are killed, we hypothesized that some CECs may induce plant cell death. Based on this hypothesis, we examined the cell death-inducing activities of expressed CECs. A previous *C. higginsianum* transcriptome study reported that *ChCEC2-1*, *ChCEC2-2*, *ChCEC3*, and *ChCEC6* were up-regulated during infection (O’Connell et al., 2012) (Supplementary Figure 2). We amplified cDNAs of *ChCEC2-1*, *ChCEC2-2*, and *ChCEC6* and synthesized the predicted *ChCEC3* CDS, then cloned these sequences into pSfinx-GW to further examine their roles in inducing cell death. Having verified that the cloned sequences were identical to the predicted CDSs reported in Tsushima et al., 2019b, they were transiently expressed in *N. benthamiana* leaves using *A. tumefaciens*-mediated transient transformation (agroinfiltration). This experiment showed that expression of *ChCEC3,* but not *ChCEC2-1*, *ChCEC2-2*, or *ChCEC6,* induced cell death in *N. benthamiana* leaves (Figure 2C).

### The Cell Death-Inducing Ability of CEC3 is Conserved among Four *Colletotrichum* Species

To investigate whether the function of *CEC3* genes is conserved across *Colletotrichum* species pathogenic on different host plants, we cloned the cDNAs of *CEC3* homologs from *C. higginsianum* (*ChCEC3*), *C. orbiculare* (*CoCEC3-1* and *CoCEC3-2*), *C. fructicola* (*CfCEC3-1* and *CfCEC3-2*), and *C. graminicola* (*CgCEC3*) into pGWB5 for expression under the control of the 35S CaMV promoter with a C-terminal GFP-tag (Supplementary Figure 3). *ChCEC3*, *CoCEC3-1*, *CoCEC3-2*, *CfCEC3-1*, and *CfCEC3-2* were identical to the previously predicted CDSs (Tsushima et al., 2019b; Gan et al., 2019 and 2020). However, *CgCEC3* from *C. graminicola* MAFF 244463 had a 30 bp insertion encoding 10 extra amino acid sequences, and a missense mutation (Supplementary Figure 4) compared to the predicted CDS of *C. graminicola* M1.001 (XM_008096207.1) (O’Connell et al. 2012) (Supplementary Figure 4A). The sequence of *CoCEC3-2* is identical to the predicted CDS encoding a 206-aa peptide, but we also cloned a shorter splice variant that encodes a 65-aa peptide due to an internal stop codon in the second exon (Supplementary Figure 5A). To distinguish the two variants transcribed from the *CoCEC3-2* gene, we refer to the longer variant having the predicted CDS as *CoCEC3-2.1* and the shorter variant as *CoCEC3-2.2*. The amino acid sequences of the cloned CEC3 homologs were predicted to have no similarity to known functional domains using InterProScan 5.39-77.0 database except for signal peptides and transmembrane helices (Mitchell et al., 2019) (Supplementary Figure 5B). The amino acid sequences of CoCEC3-2.1 and CoCEC3-2.2 were excluded due to their transmembrane helices. Alignment of amino acid sequences of the cloned homologs with CoCEC3-2.2 indicated that they are generally well-conserved except at the C-termini (Supplementary Figure 5C).

To assess if other CEC3 homologs also induce cell death, we performed an agroinfiltration assay. *ChCEC3-GFP*, *CoCEC3-1-GFP*, *CoCEC3-2.1-GFP*, *CfCEC3-1-GFP*, *CfCEC3-2-GFP*, but not *CoCEC3-2.2-GFP* and *CgCEC3-GFP*, induced cell death in *N. benthamiana* leaves by five days after infiltration (Figure 3). To investigate whether CEC3 proteins act in the extracellular or intracellular compartments, we also tested *CEC3-GFP* lacking the regions encoding predicted signal peptides (ΔSP). This experiment showed that cell death induced by transient expression of the truncated constructs tended to be stronger than full-length sequences (Figure 3). For example, although *CgCEC3-GFP* did not induce cell death, *CgCEC3ΔSP-GFP* induced weak cell death in *N. benthamiana* leaves. However, some GFP-tagged CEC3 proteins including CgCEC3-GFP and CgCEC3ΔSP-GFP did not appear to be expressed well because they were not detectable by immunoblotting (Supplementary Figure 6).

**FIGURE 3.**
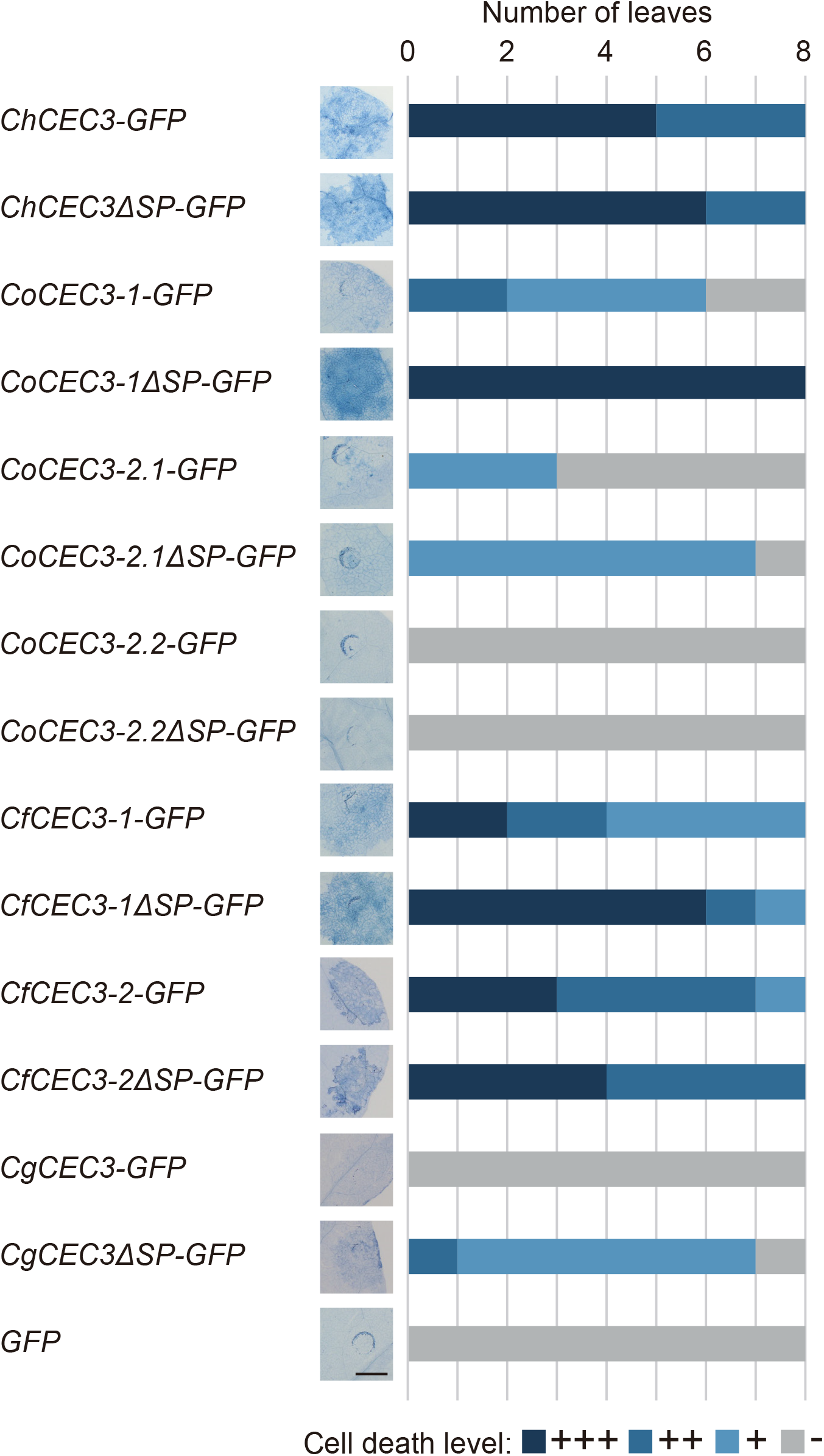
Transient expression of *GFP*-tagged *CEC3* gene-induced cell death in *N. benthamiana*.*N. benthamiana* leaves were detached five days after infiltration with *A. tumefaciens* strains carrying *GFP*-tagged *CEC3* genes in binary vectors, and stained with trypan blue to visualize cell death. Stacked bars are color-coded to show the number of each cell death level (+++, ++, +, −). Cell death induction levels were determined from observation of eight different stained leaves. Representative stained leaf images are shown on the left of the stacked bars. Bar = 5 mm.

### CEC3 Induces Plant Nuclear Expansion

To investigate CEC3 protein function in the plant cell, we observed the subcellular localization of transiently expressed ChCEC3-GFP in *N. benthamiana* leaf epidermal cells. ChCEC3-GFP was localized to mobile punctate structures, as well as the surface of spherical structures located in the center of cells expressing the protein (Supplementary Figure 7). As each ChCEC3-GFP-expressing cell always contained only one spherical structure with GFP signals at its periphery, we hypothesized that the spherical structure may be an expanded nucleus. To test this hypothesis, we transiently co-expressed ChCEC3-GFP and the endoplasmic reticulum (ER) marker HDML-mCherry (Nelson et al., 2007), which is continuous with the nuclear envelope. This experiment revealed that ChCEC3-GFP and HDML-mCherry were colocalized, indicating that ChCEC3-GFP localizes to the ER (Supplementary Figure 7). We also stained ChCEC3-GFP-expressing cells with DAPI, which showed that the spherical structures in cells expressing ChCEC3-GFP were expanded nuclei (Figure 4A). Median nuclear diameters were greater in ChCEC3-GFP-expressing cells than in GFP-expressing cells as controls (Figure 4B). The expanded nucleus phenotype was also observed in cells expressing other GFP-fused CEC3 homologs, suggesting that the function of CEC3 is conserved among homologs (Supplementary Figure 8). The detection of GFP signals on the surface of nuclei was signal peptide-dependent, as deletion of signal peptides resulted in nucleocytoplasmic localization (Figure 4A and Supplementary Figure 8). However, nuclei were still enlarged, suggesting that the signal peptide-deleted versions of CEC3 homologs also induce structural changes in the nuclei (Figure 4A, B and Supplementary Figure 8). We did not detect the nuclear expansion phenotype in CoCEC3-2.2-GFP, CoCEC3-2.2ΔSP-GFP, or CgCEC3-GFP-expressing cells. This is likely due to the low expression or instability of the fusion protein as shown in the immunoblotting (Supplementary Figure 6).

**FIGURE 4.**
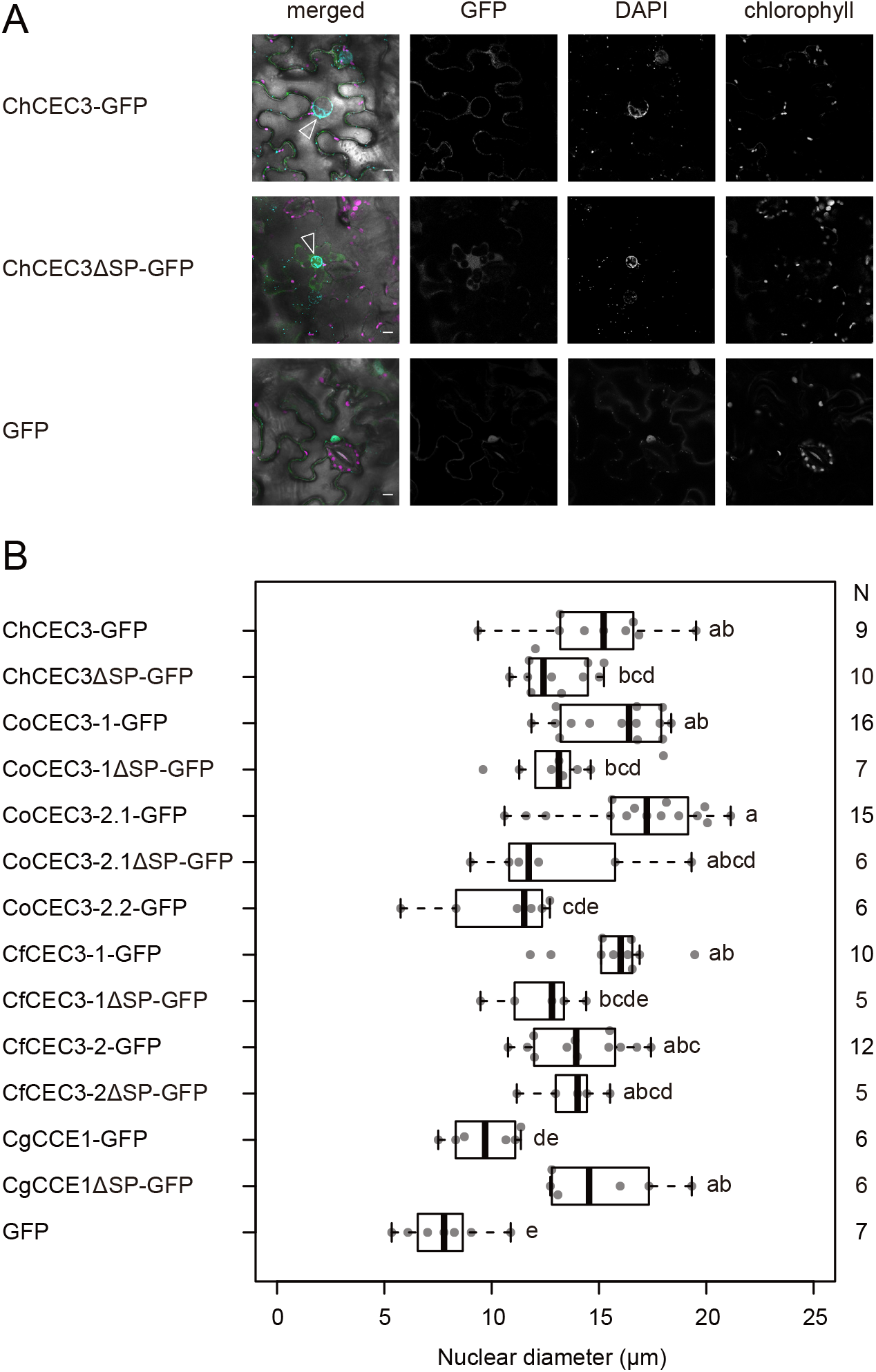
**(A)** Transient expression of GFP-tagged ChCEC3 protein-induced nuclear expansion in *N. benthamiana* leaf cells. In merged images, green represents GFP signals, cyan represents DAPI signals, and magenta represents chlorophyll autofluorescence. Open arrowheads indicate expanded nuclei. Images were taken 24 hours after infiltration. Bars = 10 μm. **(B)** Boxplots of nuclear diameters resulting from transient expression of GFP-tagged CEC3 proteins. Data represent the medians of biological replicates. N represents the number of nuclei examined. CoCEC3-2.2ΔSP-GFP is not included because no GFP signal was detected. Analysis of variance with Tukey post-hoc honestly significant difference test (*P <* 0.05) was performed.

### ChCEC3 Does Not Significantly Affect *C. higginsianum* or *Pto* DC3000 Virulence on *A. thaliana*

To investigate the contribution of *ChCEC3* to fungal virulence, we quantified its transcript levels during infection using RT-qPCR. In *A. thaliana* ecotype Col-0 leaves, the expression of *ChCEC3* was induced *in planta*, especially at 22 and 40 hours after inoculation, which correspond to the penetration and biotrophic stages, respectively (Figure 5A and Supplementary Figure 9). Next, we generated overexpression lines (Supplementary Figure 10) and knockouts in *C. higginsianum*, and inoculated *A. thaliana* ecotype Col-0 with these transformants. The lesion sizes of *ChCEC3* overexpression and knockout transformants did not differ significantly from those of the wild type *C. higginsianum* six days after inoculation (Figure 5B, C).

**FIGURE 5.**
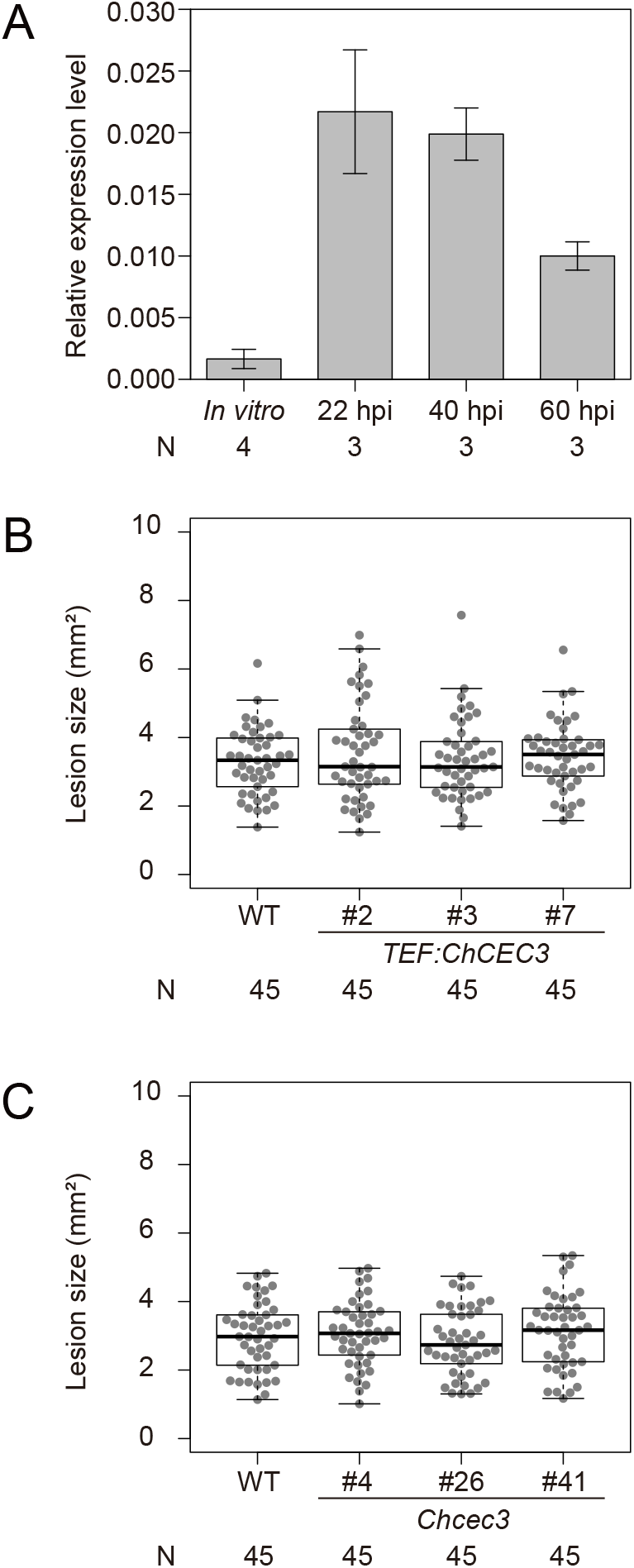
**(A)** RT-qPCR analysis of *ChCEC3* transcript levels in hyphae *in vitro* versus *in planta* infection time course. *ChCEC3* transcript levels were normalized against *ChTubulin*. Data represent the means of biological replicates. Error bars indicate standard error of the mean. N represents the number of biological replicates. **(B and C)** Boxplots of lesion area assays using *ChCEC3* overexpressors and knock-out mutants. For lesion area assays, *A. thaliana* ecotype Col-0 was inoculated with *C. higginsianum* strains. Symptoms were observed six days after inoculation. Data represent the medians of biological replicates. N represents the number of biological replicates. Analysis of variance with Tukey post-hoc honestly significant difference test (*P* < 0.05) was performed. The experiments were repeated three times with similar results.

We also evaluated the virulence effect of CEC3 on infection by the model bacterial pathogen, *Pto* DC3000 that was modified to deliver ChCEC3ΔSP, or YFP control, into plant cells using a bacterial type III secretion system-based effector delivery system with the AvrRPS4 N-terminal domain (Sohn et al., 2007). This experiment was done to test if ChCEC3 targets a general component(s) of host immunity. Expression of AvrRPS4N-HA-YFP and AvrRPS4N-HA-ChCEC3ΔSP by *Pto* DC3000 was confirmed by immunoblotting (Supplementary Figure 11A). However, the colonization of *Pto* DC3000-expressing AvrRPS4N-HA-ChCEC3ΔSP was not significantly different than *Pto* DC3000 expressing the control, AvrRPS4N-HA-YFP, at four days after infiltration (Supplementary Figure 11B).

## Discussion

Effectors play critical roles during infection by acting at the interface between microbes and host plants. Identification of conserved effectors among *Colletotrichum* fungi is thus expected to provide new insights into common infection strategies employed by members of this genus. Here, we identified effector candidates that are conserved across the *Colletotrichum* genus by performing comparative genomic analyses and showed that one of the candidates, CEC3, induces nuclear expansion and cell death in the plant.

In this study, we employed the clustering-based orthogroup inference method to identify the conservation patterns of proteins while considering evolutionary distances of 24 ascomycete fungi. Using this method, we identified seven CEC proteins that are specifically conserved in seven *Colletotrichum* species. BLASTP analysis of a wider range of organisms revealed that CEC2, CEC3, and CEC6 homologs are conserved in the genus *Colletotrichum* as well as in other closely-related fungal genera including *Pseudocercospora*, *Venturia*, *Bipolaris*, *Alternaria*, *Diaporthe*, and *Neonectria*, all of which are plant pathogens (Condon et al., 2013; Gómez-Cortecero et al., 2015; Baroncelli et al., 2016b; Chang et al., 2016; Passey et al., 2018; Armitage et al., 2020), suggesting that these CEC proteins may function as effectors in other plant-fungal pathogen interactions. BLASTP analyses showed that CEC1, CEC4, and CEC7 had limited amino acid sequence similarity despite being classified into separate orthogroups. These orthogroups may therefore have evolved different functions since their divergence, and may be worth further scrutiny. As the quantity and quality of genomic information are crucial for identifying specific/conserved genes using bioinformatic approaches, it would be of considerable interest to reanalyze the conservation patterns of effector candidates when a greater number of contiguous genome assemblies become available.

Agroinfiltration assays showed that the cell death-inducing ability of CEC3 proteins is conserved across four *Colletotrichum* species with different host specificities. Many *Colletotrichum* species, including *C. higginsianum*, establish hemibiotrophic infections, comprising an initial biotrophic phase during which they maintain host cell viability, and a later necrotrophic phase in which they elicit host cell death. The RT-qPCR experiment showed that *ChCEC3* is highly expressed in the biotrophic phase, thus CEC3 proteins may contribute to the shift in infection phases and promote colonization by initiating cell death. Alternatively, CEC3 may be recognized by a nucleotide-binding domain and leucine-rich repeat (NLR) receptor, thus leading to hypersensitive response (HR) cell death, which often limits pathogen growth (Jones et al., 2016). However, *C. orbiculare,* which expresses *CoCEC3-1* and *CoCEC3-2.1* is fully virulent on *N. benthamiana* (Shen et al., 2001), suggesting that CEC3-induced cell death is not linked to host resistance *per se*. One possible explanation, in this case, is that CEC3-induced cell death might be inhibited by other effectors such as ChEC3, ChEC3a, ChEC5, ChEC6, and ChEC34 from *C. higginsianum* and CoDN3 from *C. orbiculare* with the result that cell death is suppressed or delayed in *N. benthamiana* leaves (Kleemann et al., 2012; Yoshino et al., 2012).

Our experiments show that CEC3 proteins are likely cytoplasmic effectors because the nucleus-expanding and cell death-inducing abilities of the homologs are not eliminated by deleting the signal peptide region. In contrast, previously characterized *Colletotrichum* effectors, CtNudix from *C. truncatum* and ChELP1 and ChELP2 from *C. higginsianum* are considered to be apoplastic effectors because transient expression of the full-length constructs of CtNudix induces cell death, but not the construct lacking the signal peptide and ChELP1 and ChELP2 are confirmed to target plant extracellular components (Bhadauria et al., 2013; Takahara et al., 2016). These findings suggest that *Colletotrichum* spp. utilize both cytoplasmic and apoplastic effectors as shown in other filamentous plant pathogens (Giraldo and Valent, 2013).

One remarkable finding from this study was that the transient expression of CEC3 proteins induces nuclear expansion in *N. benthamiana* epidermal cells. Although enlarged nuclei have been observed in *Medicago truncatula* and *Daucus carota* cells infected by the arbuscular mycorrhizal fungus *Gigaspora gigantea* (Genre et al., 2008) as well as in *A. thaliana* cells infected by the powdery mildew fungus *Golovinomyces orontii* (Chandran et al., 2010), similar phenomena have not been reported in *Colletotrichum*-infected plant cells. It is possible, given that agroinfiltration provides strong transient expression in plant cells, CEC3 proteins may function without inducing nuclear expansion at endogenous expression levels during infection. Interestingly, Robin et al. reported that transient expression of the effector candidate ChEC106 from *C. higginsianum* increases nuclear areas in *N. benthamiana* epidermal cells nearly three-fold (Robin et al. 2018). While both CEC3 and ChEC106 enlarge nuclei, there are differences in their phenotypes; (i) the GFP-tagged CEC3ΔSP series are localized in the nucleocytoplasm, but GFP-tagged ChEC106 lacking the signal peptide is localized inside nuclei. (ii) The nuclei in CEC3-expressing cells are weakly stained by DAPI, but nuclei in ChEC106-expressing cells are strongly stained. (iii) CEC3-induced nuclear expansion always correlates with the cell death induction phenotype, whereas ChEC106 does not induce cell death. Thus, it is tempting to speculate that *Colletotrichum* fungi manipulate host nuclei using multiple effectors with different mechanisms of action. To our knowledge, CEC3 is the first effector candidate that induces both nuclear expansion and cell death in plants in transient expression assays. Some previous studies reported analogous enlarged nuclei during cell death in *A. thaliana* cells immediately after wounding (Cutler and Somerville, 2005) and in *Lolium temulentum* and *Sorghum bicolor* young silica cells, which deposit solid silica followed by cell death (Lawton, 1980; Kumar and Elbaum, 2018). However, the molecular mechanisms underlying these phenomena remain elusive. To dissect the nuclear expansion mechanisms and their link with cell death induction, the host target of CEC3, as well as factors involving nuclear structural changes during cell death in general, should be a focus of future studies.

The evolutionary as well as functional conservation of CEC3 proteins suggests that *Colletotrichum* spp. may target a conserved host element that is essential for plant immunity. We showed that CgCEC3 induced neither cell death nor nuclear expansion and that CgCEC3ΔSP induced weaker cell death, but did cause nuclear expansion at a similar level as other homologs. Given that *CgCEC3* was cloned from *C. graminicola*, the only monocot-infecting pathogen included in this study, CgCEC3 might be unstable, incorrectly folded, or otherwise not fully functional in *N. benthamiana*, which is highly diverged from maize, the host plant of *C. graminicola*. CEC3 proteins may have been adapted to target protein homologs in different host plants as shown in EPIC1 and PmEPIC1 from *Phytophthora infestans* and *P. mirabilis* that specialized to inhibit homologous proteases from their respective *Solanum* and *Mirabilis* hosts (Dong et al., 2014).

No virulence function for ChCEC3 was detected using fungal and bacterial systems under the conditions of this study, suggesting that ChCEC3 may be functionally redundant to other effectors in terms of its contribution to virulence, or that its contribution is minor. Plant-pathogen interactions exert strong directional selection pressure on both host and parasite, especially on genes encoding immune receptors and effectors (Plissonneau et al., 2017). Conversely, effectors that contribute little to virulence would not be subject to positive selection, and could thus remain relatively unchanged over evolutionary time. Therefore, *Colletotrichum* fungi may deploy a layer of effectors with restricted virulence effects and with functional redundancy that would result in weaker selection pressure. For a better understanding of the collective virulence effect of core effectors, further work is required, for example by using multiple knock-out mutants with a selection marker recycling system (Kumakura et al., 2019).

In this work we identified CEC3 as a highly conserved effector candidate among several other candidates in the phytopathogenic genus *Colletotrichum*. A series of analyses suggest that CEC3 proteins may have a role in manipulating host nuclei and promoting host cell death during infection. CEC3 proteins therefore could represent a novel class of core effectors that shows functional conservation in the *Colletotrichum* genus.

## Author Contribution

YT, YN, and KS: conceived the study. AT and PG: performed computational analyses and interpreted the data. AT, MN, PG, NKumakura, RH, NKato, and ST: performed the molecular biological experiments and interpreted the data. AT and PG: performed imaging analyses and interpreted the data. AT and NKumakura: prepared the figures and tables. AT, PG, NKumakura, and KS: wrote and revised the manuscript. All authors helped to edit the manuscript and approved the final version.

## Acknowledgments

We thank Ms. Akiko Ueno for technical support. We also thank Dr. Hidenori Matsui and Dr. Akira Iwase for kindly providing *A. tumefaciens* strain C58C1 pCH32 harboring pBCKH 35S promoter::GFP.

---

**Supplementary Figure 1.** | Plasmid constructions for infection assays. **(A)** pAGM4723. **(B)** pAGM4723_TEF_GFP_scd1_HygR was generated using Golden Gate cloning. **(C)** pAGM4723_TEF_ChCEC3g_scd1_HygR for generating *ChCEC3* overexpressors in *C. higginsianum*. **(D)** pAGM4723-ChCEC3KO for generating *Chcec3* knock-out mutants in *C. higginsianum*.

**Supplementary Figure 2.** | *ChCEC* transcript levels in *C. higginsianum* IMI 349063. The data are from O’Connell et al. 2012. VA: *in vitro* appressoria (22 hours post-inoculation (hpi)), PA: *in planta* appressoria (22 hpi), BP: biotrophic phase (40 hpi), NP: necrotrophic phase (60 hpi). Red letters indicate effector candidates that are up-regulated during infection. *C. higginsianum* IMI 349063 transcriptome data reported in Dallery et al. 2017 was not used because the annotation used in that study lacked the gene model for *ChCEC2-1* (CH063_14294).

**Supplementary Figure 3.** | Maximum likelihood phylogeny of CEC3 proteins. Values at nodes are based on 1,000 bootstrap replicates. Red boxes indicate CEC3 homologs cloned into pGWB5.

**Supplementary Figure 4.** | Alignments of deposited and cloned *CgCEC3* sequences. **(A)** Nucleotide sequence alignments of XM_008096207.1 and the cloned *CgCEC3* cDNA sequence. **(B)** Amino acid sequence alignments of XP_008094398.1 and the cloned CgCEC3 translated sequence. Red boxes indicate sites with differences.

**Supplementary Figure 5.** | Cloned *CEC* homologs. **(A)** mRNA structures of the cloned *CEC3* homologs. **(B)** Predicted functional domains of the cloned CEC homologs. **(C)** Amino acid sequence alignments of the cloned CEC3 proteins except CoCEC3-2.2. The sequence highlighted by a red box indicates the predicted signal peptides.

**Supplementary Figure 6.** | Immunoblotting of GFP-tagged CEC3 proteins transiently expressed in *N. benthamiana*. Samples were collected three days after infiltration. Red letters indicate lanes with bands at the expected size. Stars represent the expected sizes after signal peptide cleavage. CBB staining shows Rubisco large subunit protein as a loading control.

**Supplementary Figure 7.** | Subcellular localization of ChCEC3-GFP and ChCEC3ΔSP-GFP. *N. benthamiana* leaves were co-infiltrated with *A. tumefaciens* carrying ChCEC3-GFP, ChCEC3ΔSP-GFP, or GFP and HDEL-mCherry. Open arrowheads indicate mobile punctate structures. Images were taken at 36 hours after infiltration. Bars = 10 μm.

**Supplementary Figure 8.** | Transient expression of GFP-tagged CEC3 protein-induced nuclear expansion in *N. benthamiana* leaf cells. In merged images, green represents GFP signals, cyan represents DAPI signals, and magenta represents chlorophyll autofluorescence. Open arrowheads indicate expanded nuclei. CoCEC3-2.2ΔSP-GFP is not included because no GFP signal was detected. Images were taken 24 hours after infiltration. Bars = 10 μm.

**Supplementary Figure 9.** | Infection of *A. thaliana* ecotype Col-0 with *C. higginsianum*. **(A)** Appressoria formed on the leaf surface at 22 hours after inoculation. **(B)** An appressorium penetrating an epidermal cell to develop a small primary hypha (black arrowhead). **(C)** An appressorium forming a primary hypha (black arrowhead) and an appressorium forming an infection vesicle (white arrowhead). **(D)** Secondary hyphae (black arrowheads) growing in an epidermal cell resulting in cell death 60 hours after inoculation. Bars = 20 μm.

**Supplementary Figure 10.** | Semi-quantitative PCR analysis to confirm the constitutive expression of *ChCEC3* in fungal hyphae cultured in PD broth for two days at 24°C in the dark. *ChTubulin* was used as a reference for variation in fungal biomass.

**Supplementary Figure 11.** | Bacterial type III secretion system-based effector delivery system. **(A)** The production of AvrRPS4N-HA-YFP and AvrRPS4N-HA-ChCEC3ΔSP by *Pto* DC3000. The production of ChCEC3ΔSP was not detected due to low expression or instability of this protein. CBB staining shows Rubisco large subunit protein as a loading control. **(B)** Relative growth of *Pto* DC3000 carrying AvrRPS4N-HA-YFP and AvrRPS4N-HA-ChCEC3ΔSP in *A. thaliana* ecotype Col-0. Leaves of 5-week-old plants were hand-inoculated with OD_600_ = 0.0002 suspensions of *Pto* DC3000 strains. Samples were taken four days after inoculation to determine the extent of bacterial colonization. Error bars represent the standard deviations from the mean of eight samples for each strain. The experiments were repeated three times with similar results.

