## Supplementary Figures for "The Conserved *Colletotrichum* spp. Effector CEC3 Induces Nuclear Expansion and Cell Death in Plants"

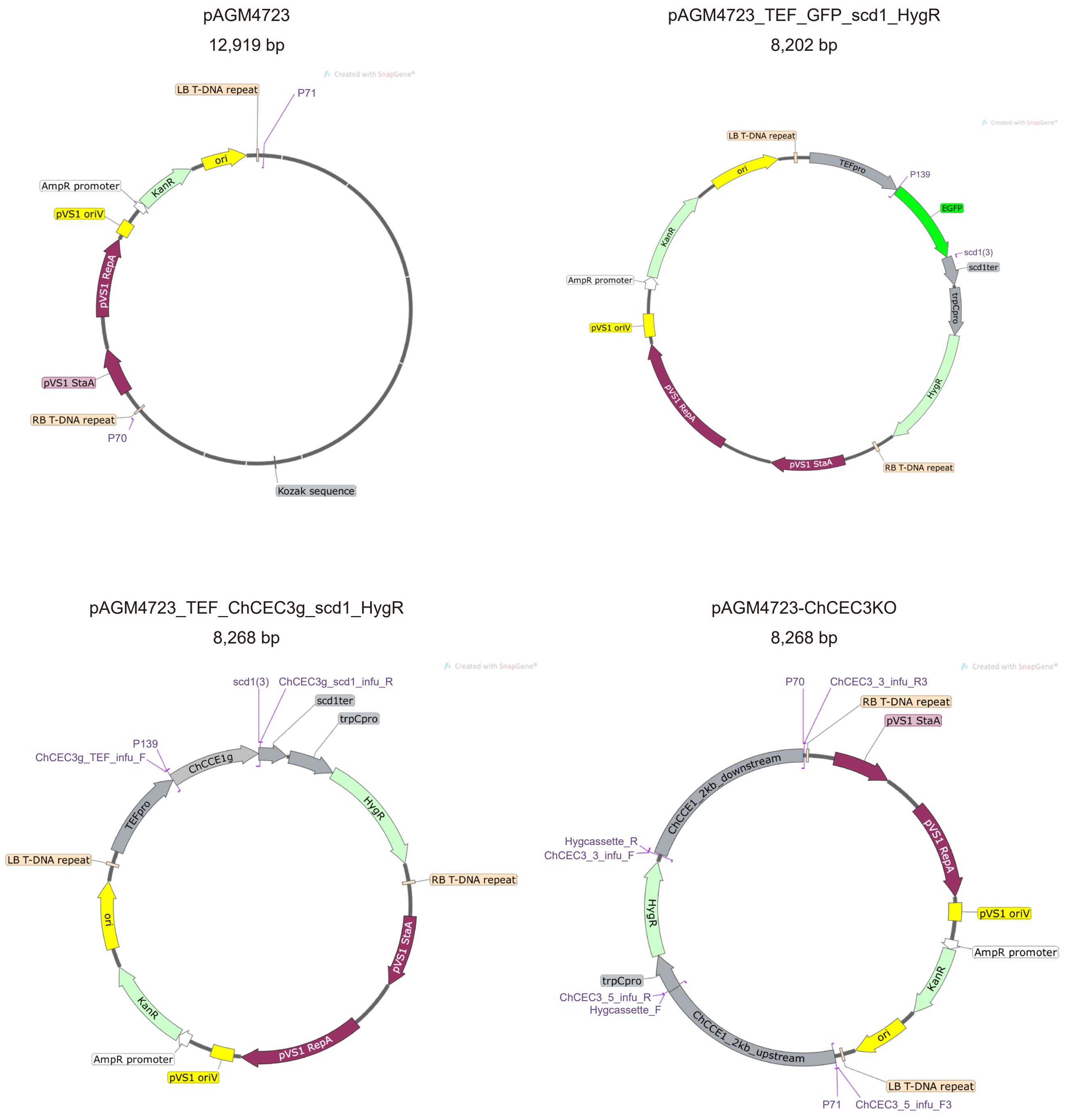


**Supplementary Figure 1.** | Plasmid constructions for infection assays. **(A)** pAGM4723. **(B)** pAGM4723 _TEF_GFP_scd1_HygR was generated using Golden Gate cloning. **(C)** pAGM4723_TEF_ChCEC3g_scd1_HygR for generating *ChCEC3* overexpressors in *C. higginsianum*. **(D)** pAGM4723-ChCEC3KO for generating *Chcec3* knock-out mutants in *C. higginsianum*.

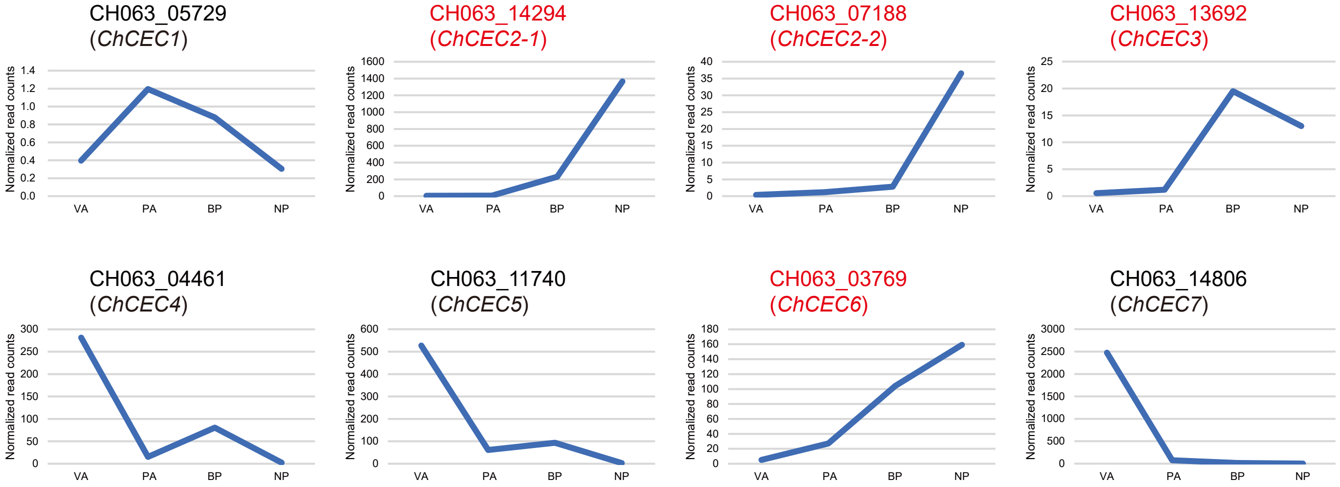


**Supplementary Figure 2.** | *ChCEC* transcript levels in *C. higginsianum* IMI 349063. The data are from O'  Connell et al. 2012. VA: in vitro appressoria (22 hours post-inoculation (hpi)), PA: in planta appressoria (22 hpi), BP: biotrophic phase (40 hpi), NP: necrotrophic phase (60 hpi). Red letters indicate effector candidates that are up-regulated during infection. *C. higginsianum* IMI 349063 transcriptome data reported in Dallery et al. 2017 was not used because the annotation used in that study lacked the gene model for *ChCEC2-1* (CH063_14294).

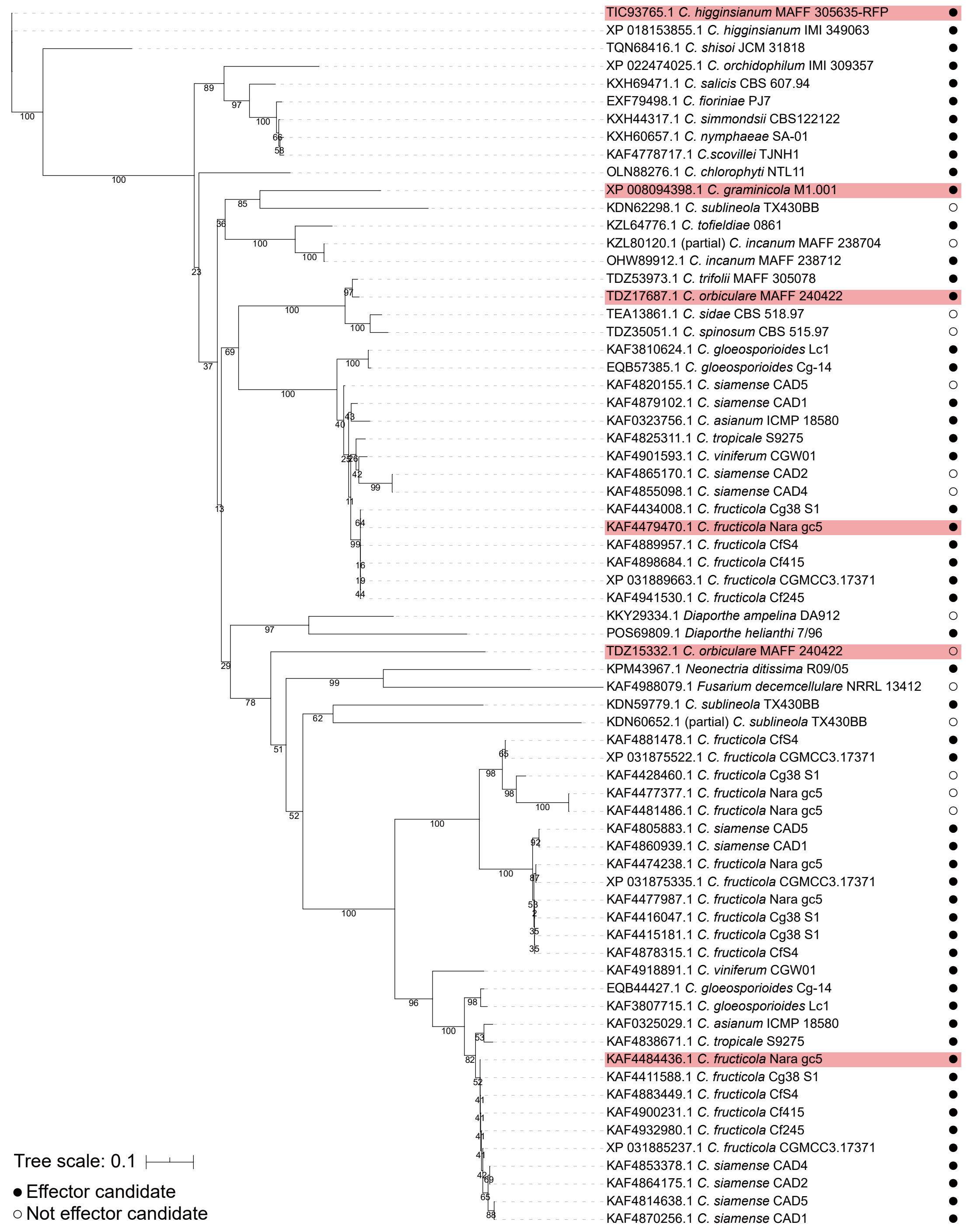


**Supplementary Figure 3.** | Maximum likelihood phylogeny of CEC3 proteins. Values at nodes are based on 1,000 bootstrap replicates. Red boxes indicate CEC3 homologs cloned into pGWB5.

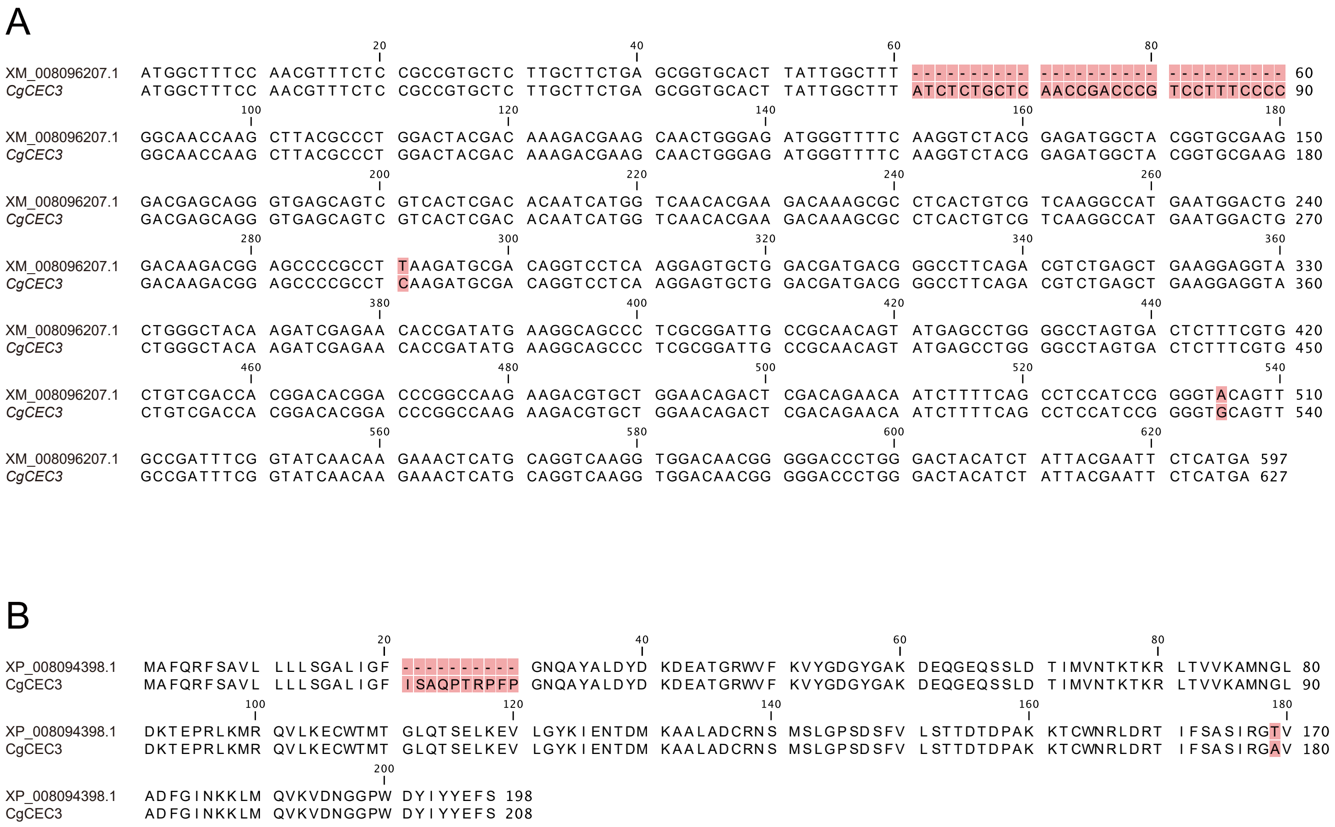


**Supplementary Figure 4. |** Alignments of deposited and cloned *CgCEC3* sequences. **(A)** Nucleotide sequence alignments of XM_008096207.1 and the cloned *CgCEC3* cDNA sequence. **(B)** Amino acid sequence alignments of XP_008094398.1 and the cloned CgCEC3 translated sequence. Red boxes indicate sites with differences.

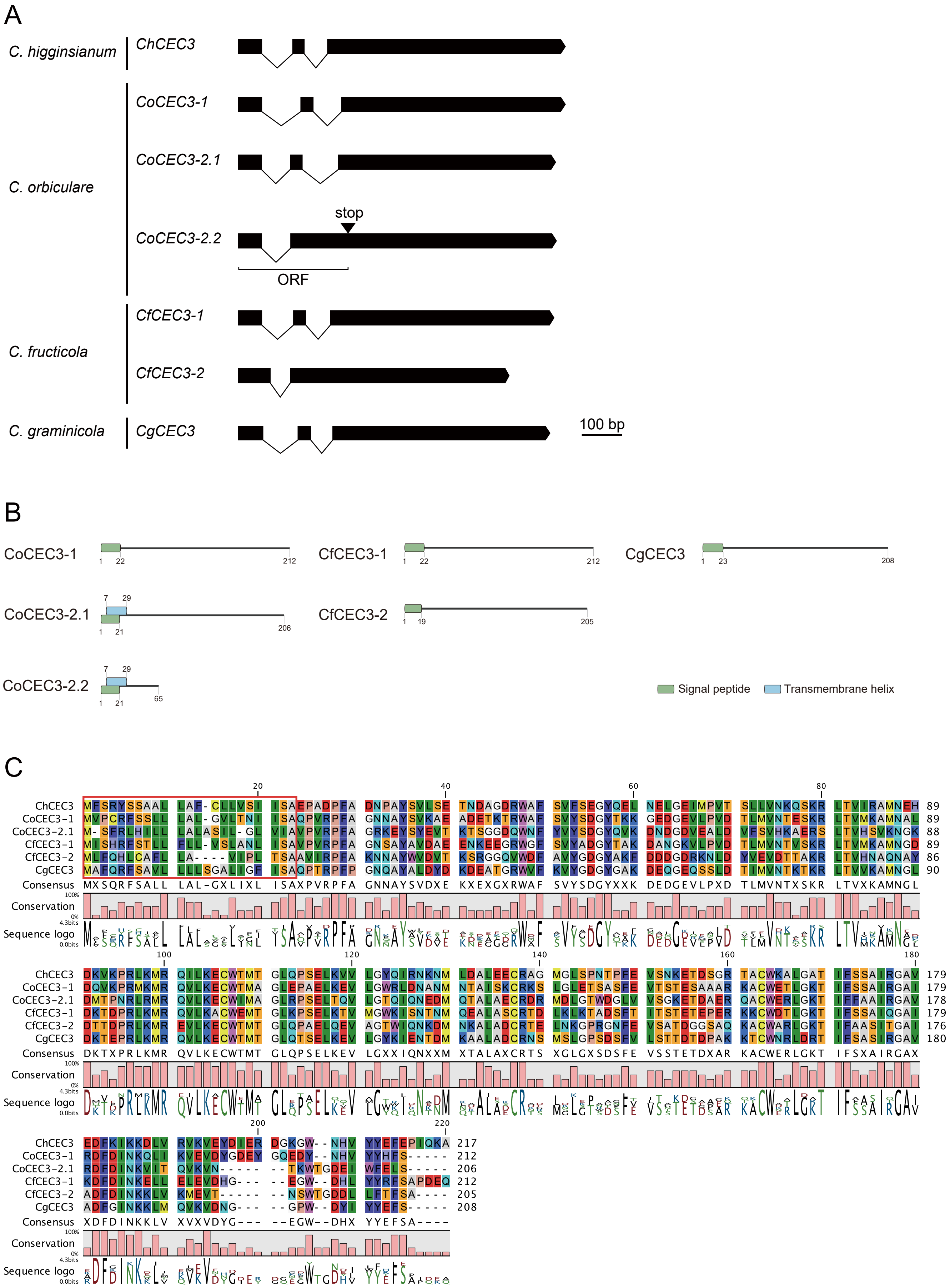


**Supplementary Figure 5.** | Cloned *CEC* homologs. **(A)** mRNA structures of the cloned *CEC3* homologs. **(B)** Predicted functional domains of the cloned CEC homologs. **(C)** Amino acid sequence alignments of the cloned CEC3 proteins except CoCEC3-2.2. The sequence highlighted by a red box indicates the predicted signal peptides.

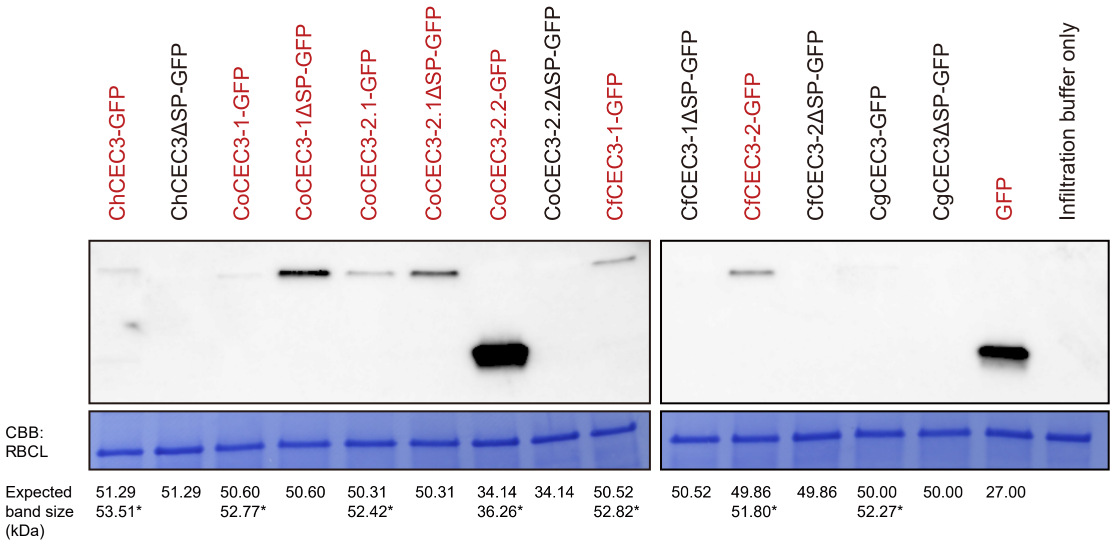


**Supplementary Figure 6. |** Immunoblotting of GFP-tagged CEC3 proteins transiently expressed in *N. benthamiana*. Samples were collected three days after infiltration. Red letters indicate lanes with bands at the expected size. Stars represent the expected sizes after signal peptide cleavage. CBB staining shows Rubisco large subunit protein as a loading control.

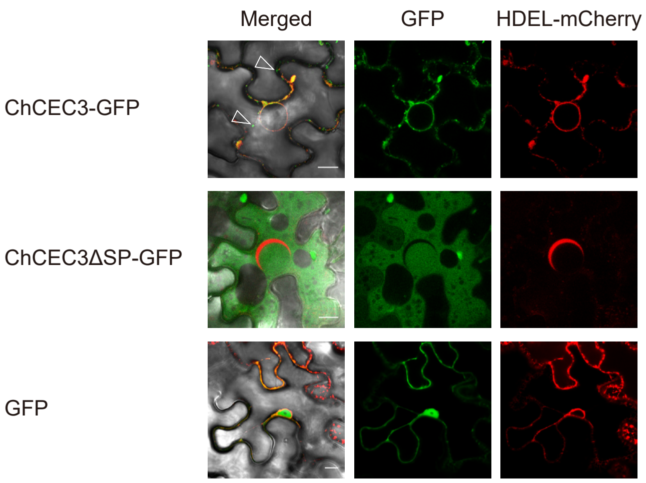


**Supplementary Figure 7. |** Subcellular localization of ChCEC3-GFP and ChCEC3ΔSP-GFP. *N. benthamiana* leaves were co-infiltrated with *A. tumefaciens* carrying ChCEC3-GFP, ChCEC3ΔSP-GFP, or GFP and HDEL-mCherry. Open arrowheads indicate mobile punctate structures. Images were taken at 36 hours after infiltration. Bars = 10 µm.

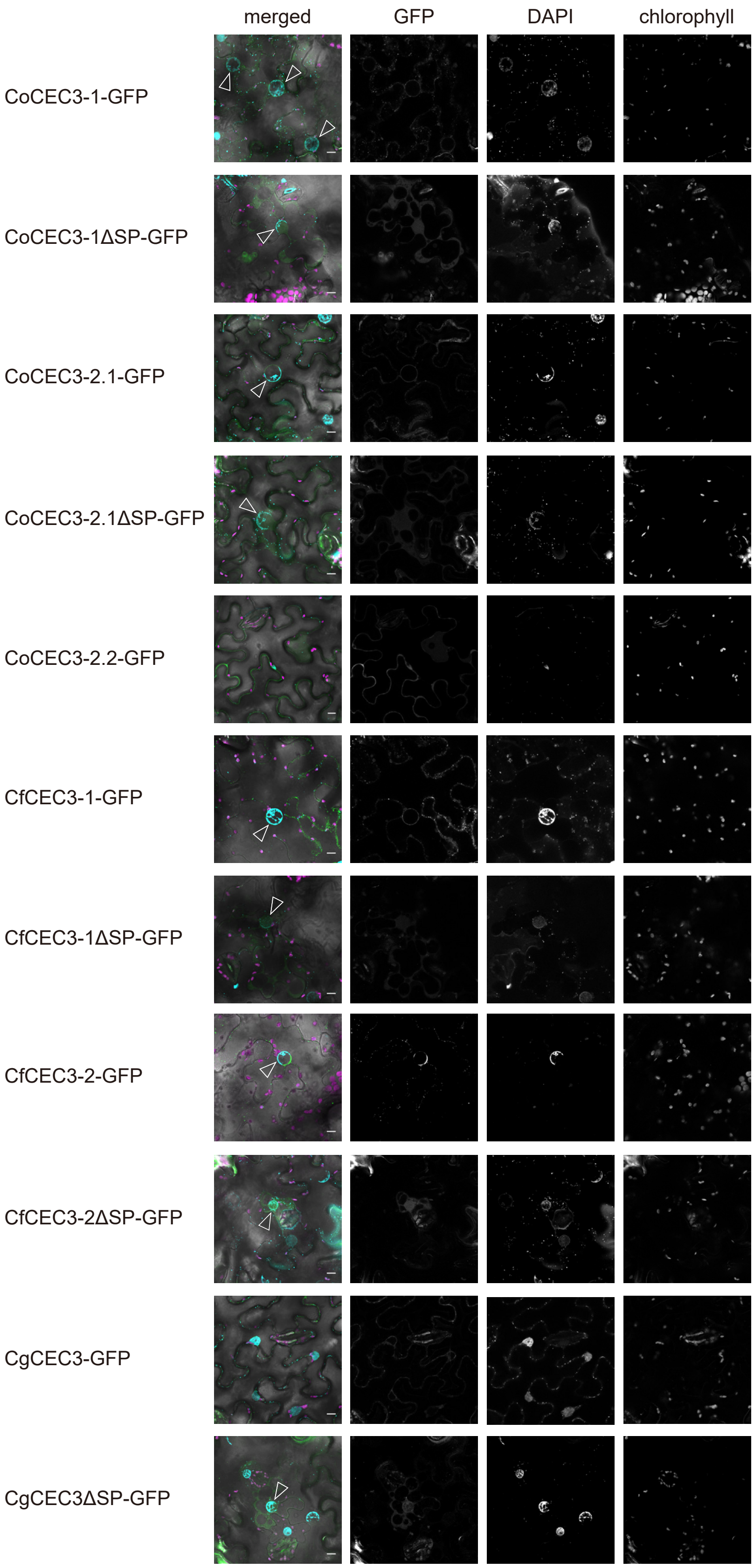


**Supplementary Figure 8. |** Transient expression of GFP-tagged CEC3 protein-induced nuclear expansion in *N. benthamiana* leaf cells. In merged images, green represents GFP signals, cyan represents DAPI signals, and magenta represents chlorophyll autofluorescence. Open arrowheads indicate expanded nuclei. CoCEC3-2.2ΔSP-GFP is not included because no GFP signal was detected. Images were taken 24 hours after infiltration. Bars = 10 μm.

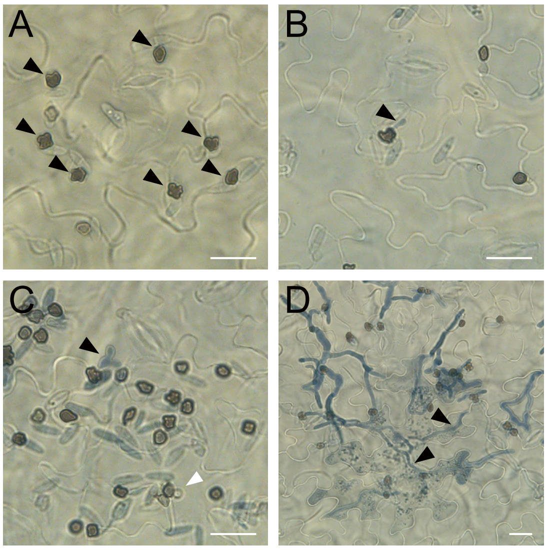


**Supplementary Figure 9. |** Infection of *A. thaliana* ecotype Col-0 with *C. higginsianum*. **(A)** Appressoria formed on the leaf surface at 22 hours after inoculation. **(B)** An appressorium penetrating an epidermal cell to develop a small primary hypha (black arrowhead). **(C)** An appressorium forming a primary hypha (black arrowhead) and an appressorium forming an infection vesicle (white arrowhead). **(D)** Secondary hyphae (black arrowheads) growing in an epidermal cell resulting in cell death 60 hours after inoculation. Bars = 20 μm.

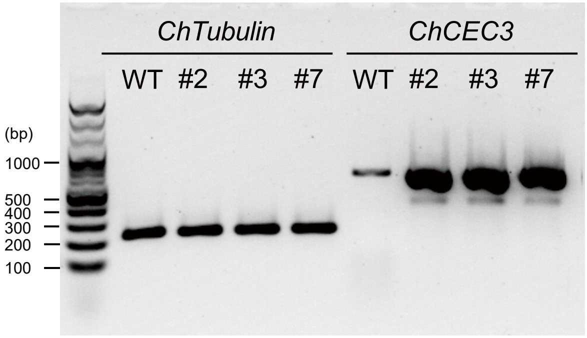


**Supplementary Figure 10. |** Semi-quantitative PCR analysis to confirm the constitutive expression of *ChCEC3* in fungal hyphae cultured in PD broth for two days at 24°C in the dark. *ChTubulin* was used as a reference for variation in fungal biomass.

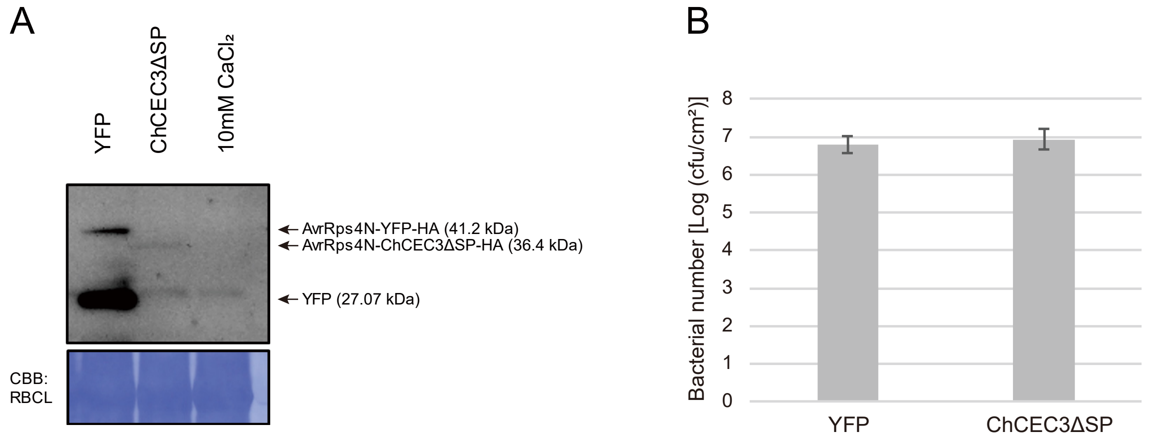


**Supplementary Figure 11. |** Bacterial type III secretion system-based effector delivery system. **(A)** The production of AvrRPS4N-HA-YFP and AvrRPS4N-HA-ChCEC3ΔSP by *Pto* DC3000. The production of ChCEC3ΔSP was not detected due to low expression or instability of this protein. CBB staining shows Rubisco large subunit protein as a loading control. **(B)** Relative growth of *Pto* DC3000 carrying AvrRPS4N-HA-YFP and AvrRPS4N-HA-ChCEC3ΔSP in *A. thaliana* ecotype Col-0. Leaves of 5-week-old plants were hand-inoculated with $OD_{600}$ = 0.0002 suspensions of *Pto* DC3000 strains. Samples were taken four days after inoculation to determine the extent of bacterial colonization. Error bars represent the standard deviations from the mean of eight samples for each strain. The experiments were repeated three times with similar results.
