## Supplementary Material 1 for "The Conserved *Colletotrichum* spp. Effector CEC3 Induces Nuclear Expansion and Cell Death in Plants"

#### Agrobacterium-mediated transformation of *Colletotrichum* fungi

1. Pre-culture
   i. Dilute 50 ul of agrobacterium stock in 5 ml of LB containing 50 µg/ml kanamycin, 25 µg/ml carbenicillin, and 50 µg/ml rifampicin in a 50 ml falcon tube. Incubate at 28˚C overnight with shaking at 220 rpm in the dark.
   Note: We use *Agrobacterium tumefaciens* strain AGL1 to transform *Colletotrichum* fungi using this protocol.
   Note: Carbenicillin and rifampicin are used to select *A. tumefaciens* strain AGL1. Kanamycin is used to select bacteria containing a binary vector.
   ii. Next morning, collect *Agrobacterium* cells by centrifuging at 4,000 x*g* for 10 min at room temperature.
   iii. Discard the supernatant and suspend the agrobacterium pellet using 5 ml of fresh Induction Medium (IM; see below). Centrifuge at 4,000 x*g* for 10 min at room temperature.
   iv. Discard the supernatant. Add 5 ml of fresh IM and check the *Agrobacterium* concentration. Dilute *Agrobacterium* to $OD_{600}$ = 0.6 in 2.0 ml of IM containing 200 µM acetosyringone and 50 µg/ml kanamycin in a 14 ml falcon tube.
   v. Incubate for 6-8 h at 25˚C with shaking at 250 rpm in the dark.

2. Prepare and dilute fungal spore suspension
   i. Make spore suspension from a sporulated plate with 0.05% Tween20.
   ii. Check spore concentration using a hemocytometer.
   iii. Dilute the suspension with 80% glycerol to give a final concentration of 15% glycerol and $10^7$-$10^8$ spore/ml. Store 100 µl aliquots at -80˚C.
   iv. Thaw the frozen aliquot at room temperature and make serial dilutions before use.
   Note: We usually prepare serial dilutions from $10^5$ spore/ml to $10^3$ spore/ml. The minimum requirement is 250 µl of $10^5$ spore/ml suspension assuming three dilutions (200 µl each of $10^5$, $10^4$, $10^3$ spore/ml).

3. Co-culture
   i. Prepare CCM plates containing 200 µM acetosyringone and 50 µg/ml kanamycin.
   Note: Use freshly prepared plates.
   ii. Carefully place 9 cm diameter Hybond -N+ membranes on CCM plates without air bubbles.
   iii. Mix 200 µl of *Agrobacterium* preculture with 200 µl fungal spore suspension.
   iv. Spread 200 µl of the culture onto each of the CCM plates covered with the Hybond -N+ membranes using sterile glass beads.
   v. Seal CCM plates using insulating tape and incubate at 25-28˚C for 24-48 h in the dark.
   Note: We usually incubate at 25˚C for 48 h.

4. Selection
   i. Transfer the Hybond -N+ membranes from CCM plates to selection plates.
   ii. Seal selection plates using insulating tape and incubate at 25-28˚C for 5-7 days.

iii. Cut about 1-2 $mm^2$ square of the Hybond -N+ membrane containing a single colony using a sterile scalpel. Transfer the membrane piece to a new selection plate.
iv. Incubate selection plates at 25˚C in the dark for a few days. After fungal colonies grow, extract genomic DNA from mycelial samples and perform genotyping.
v. Transfer the positive colonies to PDA plates and use them for experiments.

### Appendix

1. Preparation of *Agrobacterium* stocks
   i. Spread *Agrobacterium* cells harboring a binary vector on an LB plate.
   ii. Incubate plates at 28˚C for 1-2 days in the dark.
   iii. Suspend *Agrobacterium* cells in 25 ml sterile water.
   vi. Centrifuge at 4,000 x*g* for 10 min at room temperature, then discard the supernatant.
   iv. Resuspend the cells in 25 ml sterile water.
   v. Measure $OD_{600}$.
   vii. Centrifuge at 4,000 x*g* for 10 min at room temperature, then discard the supernatant.
   vi. Suspend cells in 15% glycerol to a final $OD_{600}$ = 10.
   vii. Make 50 µl of aliquots and store at -80˚C.

2. Preparation of media and reagents

- **Stock solutions**
  - 10×MM (PN): 100 mM $K_2HPO_4$ (8.71 g/500 ml), 100 mM $KH_2PO_4$ (6.81 g/500 ml),40 mM $(NH_4)_2SO_4$ (2.64 g/500 ml)
  - 10×MM (metal): 25 mM NaCl (0.73 g/500 ml), 20 mM $MgSO_4$ $7H_2O$ (2.47 g/500 ml), 7 mM $CaCl_2 \cdot 2H_2O$ (0.51 g/500 ml), 90 µM $FeSO_4 \cdot 7H_2O$ (12.5 mg/500 ml), filter-sterilized
  - 40% glucose, autoclaved
  - 80% (v/v) glycerol, autoclaved
  - AS stock: 0.2 M acetosyringone in DMSO, filter-sterilized
  - Km stock: 50 mg/ml kanamycin in water, filter-sterilized
  - Hyg stock: 50 mg/ml hygromycin B in water, filter-sterilized
  - Car stock: 50 mg/ml carbenicillin in water, filter-sterilized
  - Rif stock: 100 mg/ml rifampicin in DMSO, filter-sterilized
  - Cefo stock: 300 mg/ml cefotaxime in water, filter-sterilized
- **Minimal medium (MM)**: 10 mM $K_2HPO_4$, 10 mM $KH_2PO_4$, 4 mM $(NH_4)2SO_4$, 2.5 mM NaCl, 2 mM $MgSO_4$, 0.7 mM $CaCl_2$, 9 µM $FeSO_4$, 10 mM glucose
  i. Mix 20 ml of 10×MM (PN) and 159.1 ml of water, then autoclave.
  ii. After autoclave, add 20 ml of 10×MM (metal) and 0.9 ml of 40% glucose.
  Note: MM alone is not required for this protocol.
- **Induction medium (IM)**: MM (10 mM glucose), 40 mM MES, 0.5% glycerol, pH 5.3
  i. Mix 20 ml of 10×MM (PN) 20 ml, 158 ml of water, and MES 1.71 g.
  ii. Adjust pH to 5.3 with HCl and autoclave.
  iii. After autoclave, add 20 ml of 10×MM (metal), 0.9 ml of 40% glucose, 1.25 ml of 80% glycerol.
- **Co-cultivation medium (CCM)**: MM (5 mM glucose), 40 mM MES, 0.5% glycerol, pH 5.3
  i. Mix 40 ml of 10×MM (PN), 316 ml of water, and MES 3.41 g.
  ii. Adjust pH to 5.3 with HCl and add 6 g of Bacto agar, then autoclave.

iii. After autoclave, add 40 ml of 10×MM (metal), 0.9 ml of 40% glucose, 2.5 ml of 80% glycerol, 400 µl of Km stock, and 400 µl of AS stock.
iv. Pour the medium to sterilized Petri dishes.

- **Selection plates**

i. Mix PDA powder mix and water, then autoclave.
ii. After autoclaving, add 40% glucose, Hyg stock, and Cefo stock to a final concentration of 0.6 M glucose, 100 µg/ml hygromycin B, and 100 µg/ml cefotaxime.
Note: Cefotaxime is used to select against *Agrobacterium*.
iii. Pour the medium to sterilized 9 cm diameter Petri dishes.
